# Evolution of Human-specific Alleles Protecting Cognitive Function of Grandmothers

**DOI:** 10.1101/2021.11.26.470088

**Authors:** Sudeshna Saha, Naazneen Khan, Troy Comi, Andrea Verhagen, Aniruddha Sasmal, Sandra Diaz, Hai Yu, Xi Chen, Joshua M. Akey, Martin Frank, Pascal Gagneux, Ajit Varki

**Affiliations:** Departments of Medicine, Pathology, Anthropology and Cellular and Molecular Medicine, Center for Academic Research and Training in Anthropogeny and Glycobiology Research and Training Center, University of California San Diego, San Diego, California 92093, USA; Department of Genetics, Princeton University, Princeton, New Jersey 08544, USA; Department of Chemistry, University of California Davis, Davis, California 95616, USA; Biognos AB, Gothenburg, SE-402 74, Sweden

**Keywords:** Siglec3/CD33, pathogens, sialic acids, archaic genome, molecular dynamics simulation, phylogenetic analysis, menopause, grandmother

## Abstract

Late-onset Alzheimer’s Disease (LOAD) pathology is rare in our closest living evolutionary relatives (chimpanzees), which also express much lower microglial levels of CD33(Siglec-3)–a myelomonocytic receptor inhibiting innate immune reactivity by extracellular V-set domain recognition of sialic acid(Sia)-containing “self-associated molecular patterns” (SAMPs). We earlier showed that V-set domain-deficient *CD33*-variant allele, protective against LOAD, is derived and specific to hominin-lineage. We now report that *CD33* also harbors multiple hominin-specific V-set domain mutations and explore selection forces that may have favored such genomic changes. *N*-glycolylneuraminic acid (Neu5Gc), the preferred Sia-ligand of ancestral CD33 is absent in humans, due to hominin-specific, fixed loss-of-function mutation in CMAH, which generates CMP-Neu5Gc from its precursor, CMP-*N*-acetylneuraminic acid (Neu5Ac). Extensive mutational analysis and MD-simulations indicate that fixed change in amino acid 21 of hominin V-set domain and conformational changes related to His45 corrected for Neu5Gc-loss by switching to Neu5Ac-recognition. Considering immune-evasive “molecular mimicry” of SAMPs by pathogens, we found that human-specific pathogens *Neisseria gonorrhoeae* and Group B *Streptococcus* (affecting fertility and fetuses/neonates respectively) selectively bind huCD33 and this binding is significantly impacted by amino acid 21 modification. Alongside LOAD-protective *CD33* alleles, humans harbor additional, derived, population-universal, cognition-protective variants absent in “great ape” genomes. Interestingly, 11 of 13 SNPs in these human genes (including *CD33*), that protect the cognitive health of elderly populations, are not shared by genomes of archaic hominins: Neanderthals and Denisovans. Finally, we present a plausible evolutionary scenario to compile, correlate and comprehend existing knowledge about huCD33 evolution and suggest that grandmothering emerged in humans.

## Introduction

In keeping with the fundamental importance of reproduction for the process of biological evolution via natural selection, loss of fecundity generally coincides with the end of lifespan in almost all species studied to date. Humans and certain toothed whales like orcas are so far the only mammals known to manifest prolonged post-reproductive lifespans under naturalistic conditions [1–6]. One current explanation for such prolonged post-reproductive survival is late-life kin selection of grandmothers and other elderly caregivers of helpless young; apparently contrary to the concept of “antagonistic pleiotropy”, which posits that natural selection does not operate in late-life [7, 8]. An interesting exception is a human-specific derived allele of CD33 associated with direct or indirect protection against late-onset Alzheimer’s Disease (LOAD) [9]. Furthermore, we noted that humans harbor additional examples of such derived, population-universal gene variants that directly or indirectly impact late-life cognitive decline, which were not found in other “great ape” genomes. This was considered as genomic evidence for the evolution of human postmenopausal longevity [10]. Here we further explore the human-specific, derived alleles of genes that protect against late-life cognitive decline, and ask when and how these emerged in hominins?

In vertebrates, glycan-binding proteins of the immunoglobulin (Ig) superfamily called sialic acid (Sia)-binding Ig-like lectins (Siglecs) form a major component of the immune system [11]. As the name indicates, Siglecs recognize Sias on cell surface or secreted glycoproteins and glycolipids. Siglec-3, commonly known as CD33, is the eponymous member of the rapidly evolving subgroup of Siglecs called CD33-related Siglecs (or CD33rSiglecs) [12, 13]. In contrast, other Siglecs (Siglecs 1, 2, 4 and 15) show evolutionary conservation [14]. CD33 is a type-I transmembrane protein with an amino terminal Ig-like V-set domain followed by one Ig-like C2-set domain proximal to the transmembrane region [13]. Its cytoplasmic tail contains immunoregulatory signaling motifs called immunoreceptor tyrosine-based inhibitory motif (ITIM)s, which upon ligand binding to the extracellular V-set domain, undergo phosphorylation and recruit effector molecules like tyrosine phosphatases, SHP-1/2, which inhibit the cellular immune response. Human CD33 (huCD33) binds α2-3- and α2-6-linked *N*-acetylneuraminic acid (Neu5Ac), the predominant Sia in humans, associated either with N- and O-glycosylated molecules or sialylated glycolipids (gangliosides). HuCD33 undergoes alternative splicing, resulting in two isoforms – full length CD33M containing the ligand binding V-set domain and truncated D2-CD33 (or CD33m) lacking this domain [15]. The elimination of the terminal V-set domain is mediated through differential splicing affected by two co-inherited single nucleotide polymorphisms (SNPs) at positions rs3865444 in huCD33 promoter and rs12459419 located within exon 2 [16]. The two isoforms, CD33M and D2-CD33, differ not only in their molecular weights, but also in their cellular localization and functionality which are associated with Sia-interacting V-set domain [9, 12, 17].

HuCD33 is extensively studied for its role in different immune responses, under both normal and pathophysiological conditions including cancers [16, 18, 18–21]. Furthermore, the microglial expression of CD33 is linked with neurological pathologies like LOAD. Incidence of LOAD has been strongly associated with varied expression of CD33 isoforms in the brain of affected individuals [20, 21], where the LOAD-protective CD33 allele increases the ratio of D2-CD33 isoform relative to CD33M. CD33 is reported in almost all vertebrates, including nonhuman primates [6, 14]. While there is often high similarity in the sequence and overall genomic location, CD33 has undergone various species-specific changes. For example, murine CD33 which shows about 54% identity with huCD33 V-set and 72% identity with C2 domain, has markedly different Sia-binding and cellular expression patterns from human CD33 protein [22]. CD33 expression has greatly diverged in humans even in comparison to our closest living evolutionary relatives, the great apes. Examination of CD33 in peripheral blood showed significantly increased production of CD33M in human monocytes relative to those of chimpanzees [9]. Furthermore, the abundance of CD33 was also markedly higher in the human brain. Interestingly, although LOAD-associated neurological pathologies, for example, buildup of Aβ proteins, hyperphosphorylated tau proteins as neurofibrillary tangles, have been observed in aged nonhuman primate brains, AD has largely been regarded as a uniquely human disease [23, 24]. Interspecies variations in CD33 have also been studied in other apes like gorilla and bonobo, in comparison to huCD33 [25].

The presence of two physiologically significant isoforms, their distinct cellular localization and association with uniquely human pathologies like LOAD have made huCD33 a target of much evolutionary interest. The Sia-binding V-set domain of CD33rSiglecs including CD33 itself show high sequence variability amongst different species, often making it difficult to identify their orthologs. The selective pressure for this accelerated evolution of the V-set domains has been attributed to evasion of infectious pathogens that exploit these human innate receptors. The surfaces of each vertebrate cell are layered with tens to hundreds of million Sia-terminating glycans, forming as “self-associated molecular patterns” (SAMPs), which prevent erroneous activation of innate immune responses against the body’s own cells [26]. However, several human pathogens e.g., *Neisseria gonorrhoeae*, *Neisseria meningitidis*, *Haemophilus influenzae*, *E. coli* K1, Group B *Streptococcus*, and *Trypanosoma cruzi* cloak themselves with sialoglycans, effectively mimicking host SAMPs, and thereby avoiding the immune response [27]. Conversely, other infectious agents like influenza virus recognize SAMPs and utilize them as receptors to initiate binding and subsequent infections [28]. CD33 has also been shown to interact with Hepatitis B viral surface sialoglycans, thereby impacting its pathogenesis [29]. SAMPs and their interacting partners, Siglecs (primarily the V-set domains) are therefore continually evolving to maintain their distinctive “self-recognition” properties, while also avoiding exploitation by Sia-cloaked pathogens and parasites – a powerful example of the “Red Queen Effect” [30].

In this work, with a focus on CD33, a post-reproductive cognitive health associated human protein, we attempt to explore the evolutionary pressures that selected for unique changes in huCD33. Using human-specific pathogens like *Neisseria gonorrhoeae*, *Group B Streptococcus* and *E. coli* K1, we demonstrate differential impact of these mutations on the bacterial interactions with huCD33. We also determine the effect of these mutations on huCD33-sialoglycan binding and identify that the amino acid at position 21 within the V-set domain plays a critical role in Sia-specificity of human and chimpanzee CD33. Furthermore, we extend our study to archaic hominin genomes and show that the human-specific CD33 mutations (except the presence of truncated isoform) are shared evolutionary changes of human, Neanderthal and Denisovan common ancestor. We also expanded our analysis to include other human-specific derived genomic changes associated with cognitive health of post-reproductive human grandmothers and other elderly caregivers. Finally, we draw an evolutionary scenario to connect the current knowledge of CD33 sialoglycan recognition and pathogen engagement to propose a role for the infectious pathogens as key selective agents in human-specific CD33 evolution, generating new alleles protective against infections, that could secondarily have come under selection for their protective effects against cognitive pathologies like LOAD.

## Results

### Sequences of human CD33 extracellular domains show many changes distinct from closely related great apes

Previous investigations have identified unique properties of huCD33 that influence the functionality of this molecule in humans. The presence of a huCD33 V-set truncated isoform as well as its overall expression difference in microglia has been associated with the protection against the occurrence of neurological pathologies like LOAD in humans. Like other CD33rSiglecs, CD33 immunomodulatory roles depend both on its ligand-interacting extracellular domains and signaling motif-containing cytoplasmic tail. To gain a comprehensive understanding of different CD33 domain variations, we compared the amino acid residues of full-length CD33 from human and related nonhuman primates including chimpanzee, gorilla and bonobo (Figure 1A). While the regions encoding the C2-set domain and cytoplasmic tail are highly conserved, the amino acid residues within huCD33 V-set domain differ from their nonhuman counterparts in as many as 10 positions. Since different amino acid residues in Sia-binding V-set domain could potentially impact huCD33-sialoglycan interactions and subsequent downstream signaling pathways, we further examined the overall frequency of these changes (Figure 1B). We analyzed human sequences from the 1000 Genome database [31] and compared them with 44 gorilla, 59 chimpanzee and 10 bonobo sequences [32–34]. Most of these amino acid residues (except at positions 66 and 148) are conserved in all the great apes and appeared to have changed only in the human lineage. Interestingly, the amino acid residues at positions 66 and 148 in huCD33 are isoleucine and leucine respectively, similar to CD33 of chimpanzee and bonobo. The corresponding amino acids in its more distant evolutionary relative, gorilla, are phenylalanine (Phe) (at position 66) and valine (at position 148). The presence of the same amino acid in human, chimpanzee and bonobo at these positions suggests a more ancient occurrence of these two changes, possibly prior to the divergence of chimpanzee about 6-8 million years ago (mya). Previously it has been shown that the two linked SNPs, resulting in the splicing of the V-set truncated isoform represent a derived evolutionary modification of the CD33 proteins in humans and are absent in chimpanzees [9].

**Figure 1:**
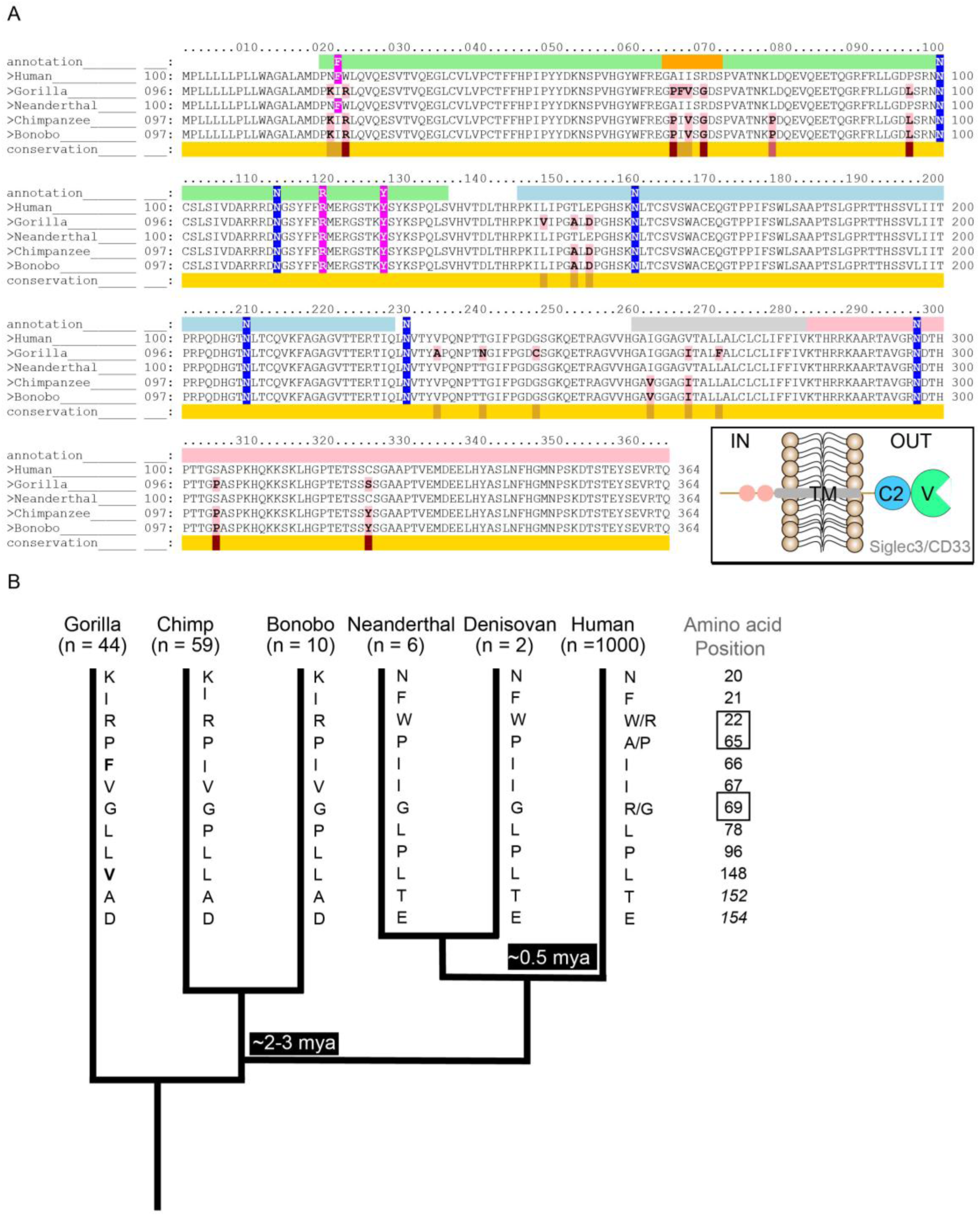
Human specific changes in CD33 are primarily present in the Sia binding V-set domain. **(A)** Comparison of amino acid sequences of CD33 from humans and “great apes” was performed using Conformational Analysis Tools software. The Great ape genomes included in the analysis are gorilla, chimpanzee and bonobo and were compared against the human protein as the template. The conservation of the sequence is indicated with yellow being the most and red being the least conserved regions. Amino acids that are different from huCD33 are highlighted in pink. Amino acids encoding the different CD33 domains are indicated above the sequence with different colors corresponding to schematic in inset, namely, V-set domain in green, C2 domain in light blue, transmembrane domain in grey and cytoplasmic end in light pink. The flexible C-C’ loop is indicated in orange. Amino acids that are in contact with Neu5Ac in huCD33 are highlighted in magenta and N-glycosylation sites in blue. (**B)** Phylogenetic analysis of the evolution of the extracellular domains of CD33 proteins from human, Great apes and archaic genomes. The number of genomes (n) for each group included in the analysis is indicated. Human and great ape CD33 sequences were compared with six Neanderthal and two Denisovan genomes. Amino acid changes present in human CD33 were also present in the ancient genomes. The positions of the amino acids that are different between human and the apes are mentioned, and the identity of the amino acid present in the corresponding positions for each group is indicated by the single letter abbreviations along the branch. Amino acids at positions 152 and 154 are within the C2 domain of CD33 protein and italicized. Polymorphisms within the human population at positions 22, 65 and 69 of CD33 protein are indicated. Amino acids in gorilla CD33 at positions 66 and 148 are different from other apes and are bold. Possible timeline for the diversion of the hominin lineage is indicated in the tree. Length of the branches in the tree is not to scale. Mya = million years ago.

To further understand the selection pressure, we calculated the nonsynonymous to synonymous substitution rate ratio (omega, ω = d(N)/d(S)) for the CD33 V-set domains of human and other great apes. The omega value of CD33 V-set domain is greater than >1 (ω =1.49) which reflects V-set domain evolution under positive selection. Subsequently, we also analyzed the Ka/Ks ratios of exon 2 sequences in every species. Except for gorilla, the other two great apes (chimpanzee and bonobo) showed Ka/Ks ratios greater than one indicating that high Ka/Ks ratio of exon 2 is not an accidental event but an evolutionary phenomenon. Taken together, these results demonstrate that CD33 in humans has been rapidly evolving possibly under positive selection, distinct from its orthologs in the great apes.

### Archaic Neanderthal and Denisovan genomes share most human CD33 protein changes, except for the SNPs for the LOAD-protective allele

Divergence of humans from other ancient hominin lineage such as Neanderthals and Denisovans has been estimated to date back approximately 0.5 mya [35]. Although full length CD33 itself is an ancient molecule, we noted that the AD-protective CD33 truncated isoform is recently derived in humans, postdating our divergence from Neanderthals and Denisovans [9]. Since huCD33 extracellular domains showed high accumulation of changes compared to the great apes, we wanted to determine if these changes were present in the common ancestor of the hominin lineage. We therefore compared CD33 protein coding sequences from 6 Neanderthal and 2 Denisovan archaic genomes obtained from the Max Planck Institute for Evolutionary Anthropology [36] (http://cdna.eva.mpg.de) with the corresponding human sequences of the 1000 Genome database (Figure 1B). Interestingly, all the amino acid residues in huCD33 that are different from the great apes are present in the ancient genomes, suggesting their occurrence in a common ancestor. These observations thus suggest that the complete loss of Sia-binding V-set domain is the latest evolutionary modification of huCD33, likely succeeding the individual amino acid changes within its extracellular domain.

### A single amino acid change facilitated CD33 engagement to the uniquely human pathogen *Neisseria gonorrhoeae*

In addition to microglial expression in the brain, CD33 is also present on tissue macrophages and peripheral blood monocytes [9]. These cells are important components of innate immune responses throughout the body, including the reproductive tract. The human female reproductive tract is also a unique niche for the microbiome, which can be invaded by important pathogens like *Neisseria gonorrhoeae* (Ng). Ng is a uniquely human infectious agent, responsible for the second most prevalent, sexually transmitted infection causative for the disease gonorrhea in human populations. Gonorrhea affects both males and females and if untreated, can have detrimental effects on reproductive health [37]. We have previously shown that Ng interacts with human CD33 but not the chimpanzee ortholog [38]. The bacterium is incapable of endogenous Neu5Ac synthesis, but instead scavenges the molecule from its host [39, 40]. Once inside the female reproductive tract, Ng utilizes the host sugar nucleotide CMP-Neu5Ac from its microenvironment to transfer Neu5Ac onto its own bacterial lipooligosaccharide. Sialylated Ng then successfully interacts with several human Siglecs including 3 (CD33), 5, 9, 11, 14 and 16 [38]. However, unlike other Siglec interactions, Ng binding to CD33 appears to be entirely Sia-dependent. Interestingly, of all the *Neisseria* species currently known, only Ng and *Neisseria meningitidis* are pathogenic to humans and both are thought to be evolutionarily young compared to others [41]. Since reproductive health/success of an organism is the key determinant of Darwinian fitness, we hypothesized that highly infectious disease like gonorrhea could potentially impact the evolution of humans, mediated through binding immune modulating proteins like CD33.

To explore our hypothesis, we examined the binding of sialylated Ng to different recombinant CD33 protein mutants, each containing the two extracellular domains with an amino acid residue changed from human to chimpanzee at the corresponding positions identified in Figure 1B. Fluorescently labelled Ng was allowed to interact with human recombinant Fc-chimeric constructs of the CD33 proteins that were immobilized onto protein A-coated plates (Figure 2A). Sia-dependence of the interaction was confirmed by comparing binding with bacteria grown in presence and absence of CMP-Neu5Ac (Supplemental Figure S1A). We observed significant reduction in bacterial binding to chimpanzee CD33 (chCD33) compared to human protein containing both V- and C2-domains (Figure 2B). However, in the absence of the V-set domain in the truncated form of huCD33 (CD33m), bacterial binding was lost. Except for the residue at position 21, all the other amino acid alterations from human to chimpanzee CD33 maintained high bacterial binding. In fact, changing the amino acid residues at positions 22, 65 (of the V-set domain), 152, and 154 (of C2 domain) increased the binding significantly compared to wildtype huCD33. In contrast, mutating the amino acid at position 21 from human to chimpanzee residue completely abolished huCD33 binding of sialylated Ng. Interestingly, mutating the chimpanzee CD33 amino acid at position 21 to its corresponding human residue enabled Ng to now engage chimpanzee CD33 (Figure 2C). Considering that Ng and its closest relative meningococcus are both uniquely human pathogens thought to have evolved from commensal *Neisseria* [42], our data suggest important implications of CD33 amino acid change at position 21 on Ng-huCD33 interaction and their mutual evolution.

**Figure 2:**
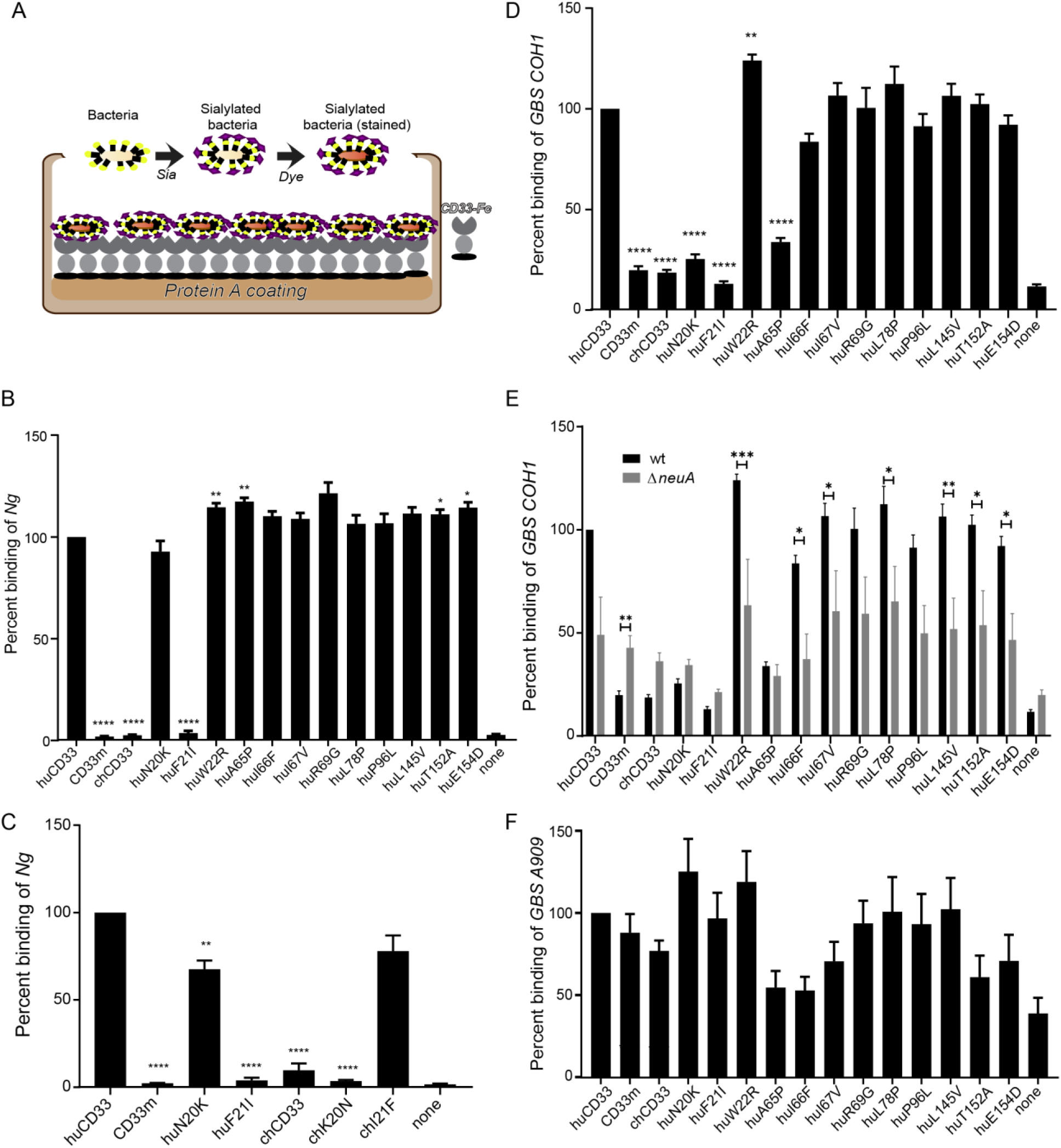
Human specific amino acid changes in CD33 affects bacterial binding. **(A)** Schematic of the ELISA-based assay using recombinant CD33-Fc chimeric proteins immobilized on protein A coated plates used to determine binding of the sialylated bacteria is shown. **(B**) Binding of fluorescently labelled *Neisseria gonorrhoeae (Ng)* was determined. The position of the amino acid different from the wildtype human CD33 protein is indicated below each bar in the x-axis. The bacterial binding to each individual CD33 mutant was normalized to the binding of wildtype human CD33 for that assay. “None” indicates no protein control for the background bacterial binding to the plate. **(B)** Binding of *Neisseria* to immobilized recombinant CD33 proteins containing the corresponding amino acid mutation (position 20 or 21) in either in human or chimp CD33 protein backbone. **(D)** Binding of Group B *Streptococcus* (GBS) COH1 strain to different CD33 mutant proteins in an ELISA based assay with immobilized recombinant CD33 proteins. **(E)** Sialic acid dependence of the binding was determined using wildtype and Δ*neuA* mutant strains of COH1. **(F)** Interaction of CD33 proteins among different GBS strains was compared using A909 and COH1 strains. ‘hu’ indicates the corresponding amino acid change in human CD33 backbone and ‘ch’ using chimp CD33. The graphs show the cumulative result from 3 independent experiments, each done in triplicate. Statistical analysis was performed in Prism software using one-way ANOVA with Durrett post comparison test. * < 0.01, ** < 0.001, *** <0.0001.

### Many amino acid changes in CD33 extracellular domains impact *GBS* engagement

While the association of *Neisseria* with CD33 is a case of Sia-mediated interaction, there are other examples of human pathogens that engage Siglecs in Sia-independent manner. One such example is Group B *Streptococcus* (GBS) which has been widely studied for its various ways of engaging host Siglecs [43]. GBS is an encapsulated pathogen commonly associated with pneumonia, sepsis and meningitis in infants and neonates. It comprises nine serologically distinct groups (Ia, Ib and II -VIII), differing in their capsular sialoglycan structures, but all containing α2-3-linked terminal Neu5Ac. Certain GBS strains have been shown to bind human Siglecs 5 and 7 in a Sia-independent manner through cell wall anchored β-protein [44], whereas some Sia-dependent binding was observed for CD33 and Siglec-9. Human Siglec-9 binding is also thought to be partially Sia-independent. Interestingly, some GBS strains are also known to interact with nonhuman primate Siglecs, for example, Siglec-9 from chimpanzee [25]. Since infections by GBS mostly impact newborns and infants, we hypothesized that it could also play a role in overall Siglec evolution in humans. Similar to the Ng-CD33 binding assay (as in Figure 2A), we examined the interactions between the recombinant CD33 proteins and GBS group III strain, COHI (Figure 2D). While the bacteria bound strongly with full-length extracellular domains of huCD33, the binding was significantly reduced in the truncated human isoform (CD33m) and the chimpanzee protein. Like Ng, GBS COHI interaction was also markedly disrupted by amino acid changes at position 21. Additionally, changing the residues at positions 20 and 65 from human to chimpanzee significantly reduced the bacterial interaction with CD33. However, GBS COHI engagement with the CD33 mutants was not entirely Sia-dependent (Figure 2E). Using GBS COHIΔ*neuA*, a mutant strain lacking its sialyltransferase enzyme (NeuA) and hence incapable of surface sialylation, we observed that about 50% of the bacterial binding to CD33 could be attributed to Sia-independent interactions. Interestingly, the CD33 binding profile of COHI was not uniform for the other serogroups of GBS, for example GBS group Ia strain, A909 (Figure 2F). None of the amino acid changes showed significant effects on CD33 interaction with GBS A909, relative to the wildtype human protein. Even the truncated human CD33 isoform (CD33m) displayed similar binding suggesting that the CD33 binding for A909 is primarily Sia-independent. Unlike Ng and GBS, we did not observe any differential sialoglycan binding with *E. coli* K1, another uniquely human pathogen of newborn infants, which contains Sia polymers on its surface (Supplemental Figure S1B). Altogether, the data demonstrate the diverse nature of CD33-interactions in three major pathogens and suggest an impact of uniquely human pathogens in the evolution of CD33 ligand-binding domain.

### Ancestral sialoglycan preference of CD33 is disrupted by amino acid change at position 21

A key change in the evolution of humans was the loss of CMP-Neu5Ac hydroxylase (CMAH), the enzyme that converts CMP-Neu5Ac to CMP-Neu5Gc resulting in a primarily Neu5Ac-rich sialome in humans, unlike any other Old-World primates, which express both Neu5Ac and Neu5Gc. This change is dated to ∼2-3 mya when human ancestors were evolving from ancestral hominins. Since we observed numerous changes mainly in huCD33 V-set domain which is critical in sialoglycan interaction and therefore important for its downstream signaling pathways, we wanted to specifically understand the effect on CD33 sialoglycan interactions. We used a microarray of chemoenzymatically synthesized glycans with defined structures, terminally capped with either Neu5Ac or Neu5Gc in different glycosidic linkages and examined their relative interactions with recombinant, soluble CD33 proteins (Figure 3). Human CD33 with V- and C2-domains bound to both Neu5Ac and Neu5Gc-terminating sialoglycans and showed maximum binding when the Sia was α2-6-linked to an underlying lactose or lactosamine glycan (Supplemental Figure S2). Most of this binding was lost in the truncated huCD33 lacking the Sia-binding V-set domain, indicating that the interactions are Sia-dependent. Conversely, the chimpanzee protein (which is identical to the bonobo orthologs and differs by only two amino acids from the gorilla) demonstrated strong preference towards Neu5Gc-terminating sialoglycans and showed almost no binding for Neu5Ac-epitopes. Considering the varied sialoglycan profiles of the two organisms, these distinct binding preferences of human and chimpanzee CD33 are interesting and suggest functional implications of the evolutionary changes in their extracellular domains.

**Figure 3:**
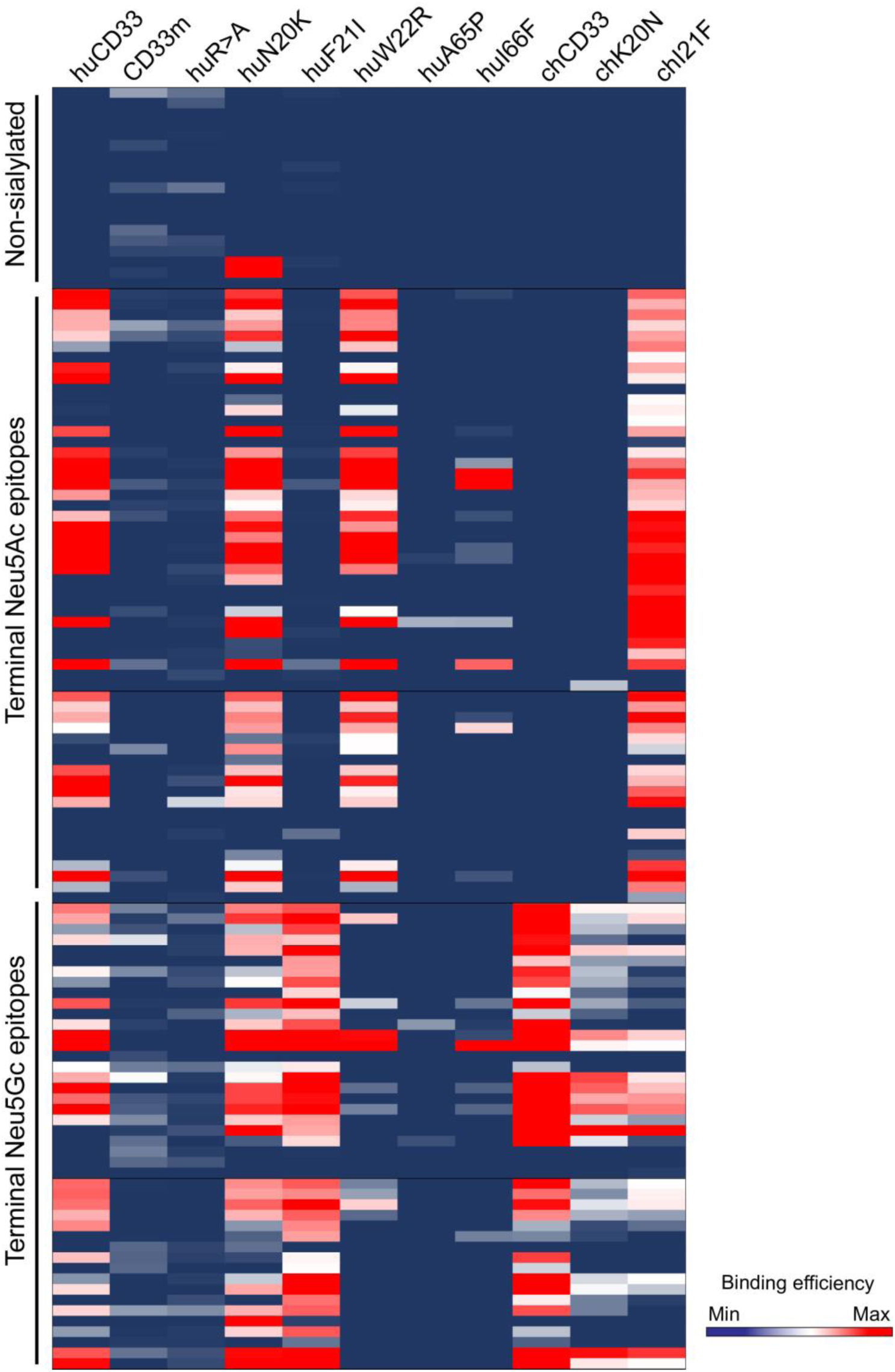
Single amino acid changes affect CD33 sialoglycan binding. Sialoglycan binding profile of purified, soluble, recombinant CD33 proteins was determined using a sialoglycan microarray containing defined, chemically synthesized glycans. Non-sialylated, Neu5Ac-and Neu5Gc-terminating glycans were grouped together in the heatmap as shown in the left. Each column indicates the binding profile of the protein indicated on the top and each row represent a distinct glycan. Blue indicates no binding and red indicates very strong binding preferences characterized by an average relative fluorescence unit (RFU) of more than 90th percentile. The result of the heatmap is summarized in Supplemental Figure S2 and the names of the individual glycans are presented in Supplemental File S1.

Because the impact of amino acid residue at position 21 was most pronounced in both of our bacterial-CD33 binding assays (Figure 2B and 2D), we next examined the influence of this change on CD33-sialoglycan binding (Figure 3 and Supplemental Figure S2). Indeed, changing the amino acid at position 21 completely altered Sia-epitope preference of CD33 for both human and chimpanzee. The presence of human amino acid residue at position 21 enabled strong binding of Neu5Ac-epitopes by chCD33, unlike its entirely Neu5Gc-preferring wildtype counterpart. On the other hand, the chimpanzee amino acid at the same position in human CD33 abolished its Neu5Ac binding. To determine if the Sia-binding changes are specific for position 21 and not an arbitrary effect of any amino acid change in V-set domain, we also looked at the Sia-epitopes of position 20 amino acid substitutions. Unlike position 21, amino acid modifications at position 20 did not have any major impact on the Sia-binding of CD33, which maintained the overall wildtype profile. Interestingly, modifications at position 22 demonstrated Neu5Ac-prefered binding for huCD33, while chimpanzee amino acid residues at 65 and 66 of huCD33 almost abolished any sialoglycan binding. Altogether, the data emphasized the importance of different amino acid changes in huCD33 V-set domain for its sialoglycan binding and identified the amino acid at position 21 to be critical in the functionality of CD33 protein.

### MD simulations provide structural insights for the differences in Sia-binding preference between human and chimpanzee CD33

We performed an extensive theoretical investigation based on molecular dynamics simulations. A detailed analysis of several available crystal structures of huCD33 revealed that the V-set domain is dynamic. For example, the C-C’ loop as well as the side chains of phenylalanine at position 21 (Phe21) and histidine at position 45 (His45) are resolved in two different conformations in PDB entry 5ihb (Supplemental Figure S3). Of all the amino acids that differ between human and chimpanzee, only the side chain of Phe21 is in direct contact with a bound Neu5Ac residue in the crystal structures of huCD33 (through the methyl group at position 5). Based on the assumption that Neu5Gc binds to the same binding site as Neu5Ac, the change in binding preference from Neu5Gc (in chimpanzee) to Neu5Ac (in human) cannot be explained by a simple I21F mutation. Both amino acids have hydrophobic side chains that cannot establish favorable interactions with the polar glycolyl group. Consequently, there is probably a more complex reason for the shift of binding preference. Based on data derived from 47 molecular dynamics (MD) simulations covering an accumulated timescale of more than 100 µs we conclude that in chCD33 His45 adopts mainly the ‘up’ conformation (Figure 4A), which allows favorable hydrogen bonding with the glycolyl group. MD simulations (as well as x-ray crystallography) show that in huCD33 His45 can also exist in the ‘up’ conformation (Figures 4C and Supplemental Figure S4), which would be compatible with favorable Neu5Gc binding. However, when His45 is in the ‘down’ conformation Phe21 can stack partly with tyrosine (Tyr) at position 127 (Figure 4B) forming a small hydrophobic pocket, which allows the methyl group of Neu5Ac to bind favorably. To demonstrate if the binding affinity difference between Neu5Ac and Neu5Gc may be indeed correlated to the up/down conformational equilibrium of His45, we performed a series of MD simulations of chCD33 on the microsecond timescale where Neu5AcOMe or Neu5GcOMe molecules are present in the solution. The lifetimes of the complexes spontaneously formed during the MD with Neu5Gc are on average much longer when His45 is ‘up’ (Figure 4D top, Supplemental Figure S4). In contrast the lifetimes of the complexes spontaneously formed with Neu5Ac are much shorter independent of the conformational state of His45 (Figure 4D bottom), which would explain the lack of measurable binding affinity of Neu5Ac to chCD33. In summary, our extensive MD simulations - including unbiased simulation of carbohydrate binding and unbinding events - could provide a reasonable explanation for a change in binding specificity that is likely to be caused by an alteration of the protein-ligand interaction pattern remote from the mutated amino acid.

**Figure 4:**
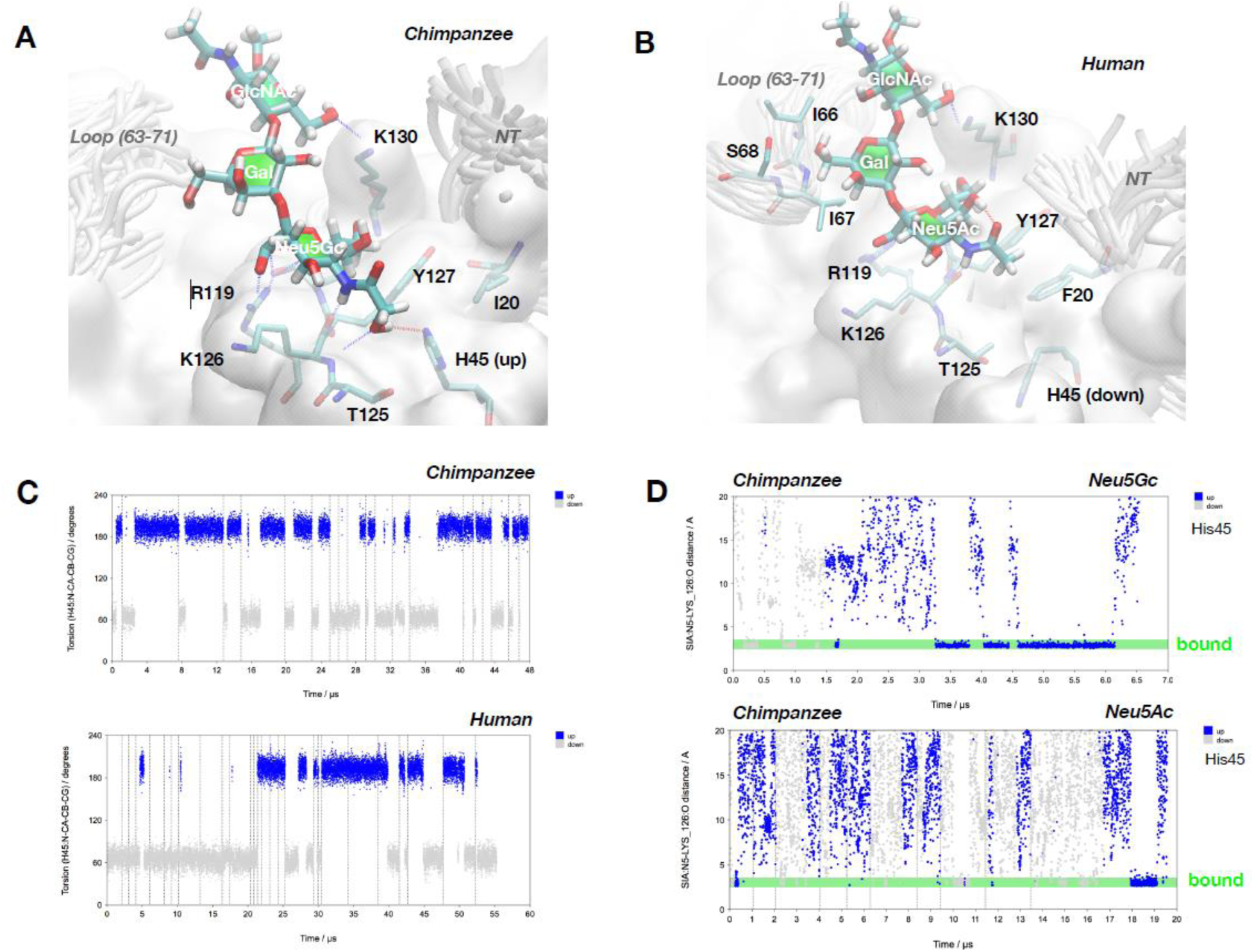
Structural modeling to understand the differential binding preference of human and chimpanzee CD33 proteins. **(A)** 3D model of the complex between Neu5**Gc**α2-3Galβ1-4GlcNAcβOMe and chCD33. The increased affinity of Neu5Gc may be explained by intermolecular hydrogen bonds involving the OH-group of Gc. It should be noted that the number of favorable interactions is maximal when His45 is in ‘up’ conformation. **(B)** 3D model of the complex between Neu5**Ac**α2-3Galβ1-4GlcNAcβOMe and huCD33. The methyl group of Ac is located in a small hydrophobic pocket formed by the side chains of Tyr127 and Phe20. It should be noted that His45 is in ‘down’ conformation because otherwise – in the conformation shown -the bulky side chain of Phe20 would overlap partly with His45 in ‘up’ conformation. **(C)** Molecular dynamics of His45 side chain orientation. Accumulated MD trajectories of torsion angle N-Cα-Cβ-Cγ are shown. The ‘up’ conformation is present when torsion values are fluctuating around 200 degrees and the ‘down’ conformation is characterized by values around 70 degrees. For chimpanzee, it can be observed reproducibly that simulations started with His45 in ‘down’ conformation undergo a transition to the ‘up’ conformation on the microsecond timescale. In contrast the ‘down’ conformation appears to be more stable in huCD33, which would make Neu5Ac binding more likely. **(D**) MD simulation of unbiased binding and unbinding events of Neu5Gc (top) and Neu5Ac (bottom) to chCD33. For Neu5Gc the lifetime of the complex is significantly longer when His45 is in ‘up’ conformation, as can be seen from the 6.5 µs MD simulation shown on the top. Also, for Neu5Ac multiple binding and unbinding events occurred on a timescale of about 20 µs, however in general (with one exception) the lifetimes of the complexes formed are significantly shorter than for Neu5Gc.

### Human-specific polymorphisms in cognitive-health related genomic variants are present in all human populations

In an earlier study we observed several genes, directly associated with neurodegenerative diseases or correlated with aggravation of the cognitive decline in aged-population, are derived alleles in humans [9]. Increasing evidence of correlation between cognitive health and non-neurological, metabolic conditions, e.g., diabetes [45, 46] suggest that such derived alleles could be important in the maintenance of cognitive health in human grandparents. Here, we expanded this list of cognition-protective gene variants through literature and database (https://alzoforum.org) searches [20, 47–58] to include additional gene variants, namely *BINI, ARID5B, PICALM, PILRA*. Supplemental Table S1 describes the characteristics of 13 human genes that are implicated in diseases including dementia, cardiovascular diseases (CVD), hypertension and AD. While some of these physiological abnormalities like salt retention, hypertension, diabetes, appear non-neurological, they have been associated with the aggravation of the pathologies resulting in late-life cognitive decline [59]. Notably, the derived alleles are common and found in globally diverse human populations, indicating that they predate the common ancestor of modern humans (Supplemental Table S2).

### SNPs associated with human-specific cognitive protective alleles are unusual in their absence in the archaic hominin genomes

With the availability of genomes from extinct archaic hominins [36, 60, 61], a set of SNPs can be assessed as to whether their protective phenotypes arose recently in the evolutionary history of anatomically modern humans. We previously showed many other human-chimpanzee differences were shared with archaic hominins (Denisovan/Neanderthal) genomes; for example, genomic changes in CD33rSiglecs [62]. To gain similar insights about the evolutionary origin of these cognitive-protective loci, we analyzed the Neanderthal and Denisovan reference genomes and compared them with modern human sequences. Analysis of the 1000 Genomes dataset shows the presence of protective alleles in human populations with variable frequency (Table 1). Analysis of the available genomic data from Neanderthal and Denisovan genomes showed that only two derived variants (rs2975760 and rs2588969; Table 1) are present in these archaic genomes, suggesting the remaining eleven derived, protective variants arose after the divergence of modern and archaic hominins approximately 0.5 mya [35, 63]. This is in striking contrast to most human-chimpanzee genomic differences in which the archaic hominins are similar to humans. In fact, majority of the Sia-related genes lack positive selection signatures and rather show neutral evolution in the modern human lineage [64]. To more formally assess whether the high frequency, global distribution, and recent origin observed for eleven of the thirteen SNPs is unusual, we performed a resampling analysis of variants in the genome. Variants in the 1000 genomes dataset that met the following criteria were considered: 1) present in both Altai Neanderthal and Denisovan minimal filters, 2) derived in at least one modern individual from non-admixed African populations, 3) called in both archaic samples, and 4) have an ancestral allele matching the reference or alternative allele. To eliminate any bias in the analysis and match the allele frequency (AF) of these SNPs compared with that of any random SNPs, we first matched our universe of SNPs to the 13 SNPs of interest by AF, ± 2 derived haplotypes (Figure 5). Resampling was then performed by drawing a SNP from each of the 13 matched sets and assessing how many derived alleles were observed, resulting in a *p*-value = 0.08333 ± 0.00003. As a less conservative estimate, directly sampling from the universe of SNPs and estimating the probability of observing at most two derived SNPs and a mean allele frequency as large as the empirical variants of interest produced a highly significant *p*-value = 0.00487 (Supplemental Figure S5). Repeating either analysis on the set of other Siglec-related SNPs indicates they are consistent with a random draw from the genome [62]. Regardless of the individual limitations, taken together our phylogenetic analyses demonstrate the unique patterns of allele frequencies in worldwide populations distribution of these thirteen late-life cognitive decline linked SNPs (Figure 5 and Supplemental Figure S5). Interestingly, co-inherited CD33 SNPs associated with the cognitive health in LOAD are present only in modern human genomes [9]. A noteworthy example in our list is the human gene encoding the protein, apolipoprotein E (APOE), involved in fat metabolism in mammals. *APOE* gene exists in three allelic variants (E2, E3 and E4) where APOE4 is associated with high risk of LOAD and other allele like APOE2 is protective against the cognitive decline in elderly caregivers [65]. Interestingly the presence of APOE4 is also correlated with the protection from severe diarrhea in children [66]. While conclusive determination of the positive selection of these alleles in modern human requires further analysis, our data suggest that the evolutionary origin of most of these cognitive-health protective changes followed the divergence of modern humans from archaic genomes. This is also supported by the presence of grandparents, uniquely in humans. Regardless, the process of evolutionary emergence of each of these alleles is likely to be distinct and deserves further investigation.

**Figure 5:**
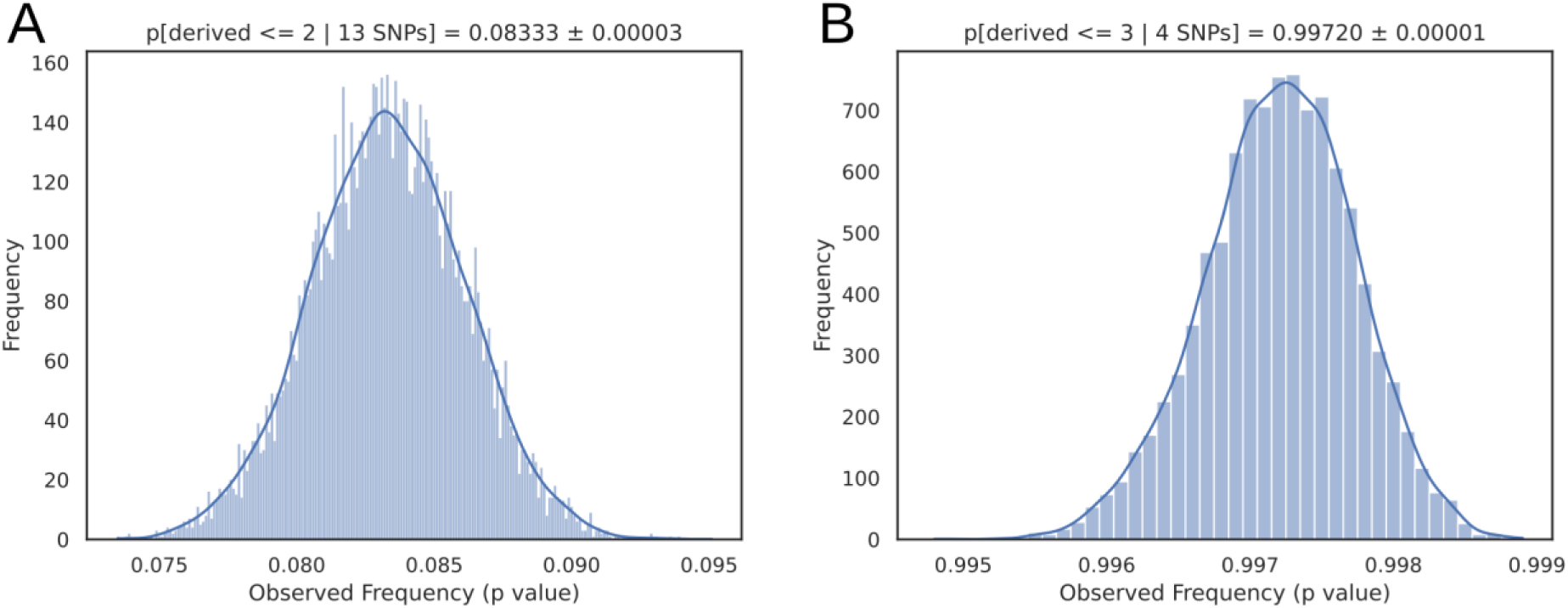
Resampling analysis with matched allele frequency SNPs from 1000 genomes variants. Frequency distribution of SNPs with similar properties to the LOAD protective set **(A)** and other Siglec SNPs **(B).**

**Table 1:** Gene variants directly or indirectly affecting cognitive function. Allele has the derived allele as the lower, bolded entry. Archaic genotypes are reported for SNPs passing all quality filters.

|  |  |  |  |  |  | Archaic Genotype |  |
| --- | --- | --- | --- | --- | --- | --- | --- |
| Gene | Associated Disease<sup>a</sup> | SNP ID | hg19 Position | Allele | Derived Global AF<sup>b</sup> | Altai | Denisovan |
| *AGT* | Sodium retention | rs699 | 1:230845794 | G | 0.295 | 0/0 | 0/0 |
|  |  |  |  | **A** |  |  |  |
| *BIN1* | AD | rs7561528 | 2:127889637 | A | 0.8 | 0/0 | 0/0 |
|  |  |  |  | **G** |  |  |  |
| *SGC2* | Hypertension | rs1017448 | 2:224466344 | T | 0.879 | 0/0 | 0/0 |
|  |  |  |  | **C** |  |  |  |
| *CAPN10* | Type II Diabetes | rs2975760 | 2:241531163 | C | 0.882 | 1/1 | 1/1 |
|  |  |  |  | **T** |  |  |  |
| *PPARG* | Type II Diabetes | rs1801282 | 3:12393125 | C | 0.070 | 0/0 | 0/0 |
|  |  |  |  | **G** |  |  |  |
| *CYP3A5* | Sodium retention | rs776746 | 7:99270539 | T | 0.621 | 0/0 | 0/0 |
|  |  |  |  | **C** |  |  |  |
| *ARID5B* | AD | rs2588969 | 10:63611354 | A | 0.532 | 0/0 | 1/1 |
|  |  |  |  | **C** |  |  |  |
| *SPON1* | Dementia | rs2618516 | 11:14021639 | C | 0.341 | 0/0 | 0/0 |
|  |  |  |  | **T** |  |  |  |
| *PICALM* | AD | rs10792832 | 11:85867875 | G | 0.313 | 0/0 | 0/0 |
|  |  |  |  | **A** |  |  |  |
|  |  | rs3851179 | 11:85868640 | C | 0.315 | 0/0 | 0/0 |
|  |  |  |  | **T** |  |  |  |
| *APOE* | LOAD | rs429358 | 19:45411941 | C | 0.849 | 0/0 | 0/0 |
|  |  |  |  | **T** |  |  |  |
|  |  | rs7412 | 19:45412079 | C | 0.075 | 0/0 | 0/0 |
|  |  |  |  | **T** |  |  |  |
| *CD33* | LOAD | rs3865444 | 19:51727962 | C | 0.211 | 0/0 | 0/0 |
|  |  |  |  | **A** |  |  |  |
| *PILRA* | AD | rs1859788 | 7:99971834 | G | 0.341 | - | - |
|  |  |  |  | **A** |  |  |  |
| *TCFLC2* | Type II Diabetes | rs7903146 | 10:114758349 | T | 0.772 | - | - |
|  |  |  |  | **C** |  |  |  |
| *CD33* | LOAD | rs12459419 | 19:51728477 | C | 0.211 | - | - |
|  |  |  |  | **T** |  |  |  |

<sup>a</sup> See supplemental Table S1 for details and the primary literature citations.
<sup>b</sup> See supplemental Table S2 for the global population distribution.

## Discussion

Fossil evidence and genomic comparisons leave little doubt about the fact that our species evolved from an African hominin. However, a detailed understanding of modern human origins is plagued by numerous uncertainties, with regard to the identity of the ancestral lineage and precise geographic locations. Evolution of modern humans was accompanied by many anatomical and behavioral changes, but increasing evidence suggests it also included uniquely human-derived modifications in the genome compared to the archaic genomes (Neanderthal/Denisovan) or the genome of the great apes [9, 62]. Taken together with our previous study [9], we have identified many such human-specific genes associated with cognitive health of grandmothers and other human elders who are often involved in the caregiving of the young. These findings, which appear paradoxical to the concept of senescence due to antagonistic pleiotropy, have lent much additional support to the “Grandmother hypothesis” [1] bolstering the case for selection of human female post-reproductive survival and the existence of grandmothers. Unlike in any other mammals (except orcas and some other toothed whales), the occurrence of this prolonged post-reproductive life span in humans has stirred scientific interest. While deciphering the precise evolutionary course of any gene/protein is challenging and the proposed schemes/players are not entirely verifiable, here we attempt to compile the current evolutionary and experimental findings of one such protein associated with late-life cognitive-decline: CD33.

A ratio of high wildtype human CD33 and a low truncated isoform of CD33 have been implicated in the progression of LOAD associated with the cognitive health of elderly population. In contrast, LOAD is unknown in chimpanzees, although evidence of LOAD pathologies has been observed in some chimpanzee brains. We found that human CD33, which is highly expressed in microglia of the human but not chimpanzee brain, recognizes Neu5Ac – the predominant Sia synthesized in humans – as self-associated molecular patterns (SAMPs). In contrast, our closest evolutionary relative, the apes and other Old-World primates contain both Neu5Ac and Neu5Gc. We found that the ancestral form of CD33 in chimpanzees and other great apes selectively recognizes only Neu5Gc-glycans as SAMPs (Figure 3). Notably, Neu5Gc – the ligand recognized by chCD33 is rare in chimpanzee brain, and there is also significantly less chCD33 protein compared to CD33 in humans [9]. On the other hand, SNPs resulting in the truncated CD33 have only been observed in the human genome and not any of the archaic or great ape genomes. We also find that the truncated human CD33 does not interact with Sia (Figure 3). Taken together, these observations suggest that full-length CD33-Sia interactions are stronger in human brain compared to chimpanzee and the human-specific SNPs in CD33 resulting in the truncated protein abolish this interaction. The question remains what could have possibly led to the selection of the truncated isoform of human CD33 that does not interact with Sia. In this regard, CD33 on macrophages plays crucial roles in different immune responses as well as during infections. Human CD33 has also recently been shown to be involved in immunomodulation during infection with hepatitis B virus [29]. Our previous and current data show that uniquely human pathogens like *Neisseria* and GBS display Neu5Ac that is recognized as ‘self’ by human but not chimp CD33 [38]. In the current work, we further found that the Sia-binding-domain-depleted, truncated human CD33 isoform doesn’t bind and thus escape exploitation by sialylated pathogens (Figure 2). This suggests that this truncated CD33 may have been an adaptation to counter the CD33-exploiting, immune-evasive behavior of pathogens like *Neisseria* and GBS.

Taking together all currently available experimental data (including this study) we attempt to draw a plausible evolutionary scenario for CD33 protein evolution in humans and present in the context of relevant evolutionary events (Figure 6). We hypothesize that the scarcity of the strongly preferred Neu5Gc ligand of ancestral CD33 in the brains of chimpanzee (and other great apes) was associated with low microglial expression. Subsequent hominin loss of CMAH (i.e., complete loss of Neu5Gc ligand) could then have selected for the upregulation of CD33 levels perhaps to compensate for the loss of ligands, a change to Neu5Ac-binding preference, and functional recruitment of CD33 to human microglia. Alongside the microglial CD33, the corresponding changes in the tissue macrophage proteins might have facilitated the emergence of Neu5Ac-coated pathogens (for example, *N. gonorrhoeae* and Group B *Streptococcus*) that evolved “molecular mimicry” of Neu5Ac-SAMP ligands to manipulate the immune response. Appearance of the truncated isoform lacking the ligand-binding domain (CD33m), then probably allowed CD33 to escape the immune evasion by these sialylated pathogens (Figure 2). This selection pressure to stop manipulation by sialylated pathogens could have also altered splicing towards a higher level of truncated CD33, which also gets diverted to peroxisomes [12]. While the significance of this diversion is unclear, decrease of full-length CD33 would facilitate escape from Neu5Ac-coated, CD33-engaging pathogens. Finally, sometime during the last 1 million years, increased brain size presumably selected for early, short interbirth interval in human, which might have resulted in more helpless young, requiring cooperative breeding and caregiving. However, the value of postmenopausal grandmothers and other elderly caregivers would then have been blunted by the appearance of LOAD. The synthesis of the truncated isoform of CD33 protects from *Neisseria* during reproductive age and a higher ratio of truncated to full-length isoforms correlates to decrease of LOAD in grandmothers. However, a small amount of the full-length isoform remains, likely to downregulate hyper-inflammation that might arise during prolonged absence of SAMP-recognition. Notably when an elderly caregiver gets LOAD, not only are the evolutionary benefits of the individual lost, but this also presents an increased burden to care for that elder individual. Altogether under this proposed scenario, the current state in the evolution of human CD33 protein represents a trade-off between the evolutionary response to exploitation by pathogens in early life and cognitive maintenance in post-reproductive late life.

**Figure 6:**
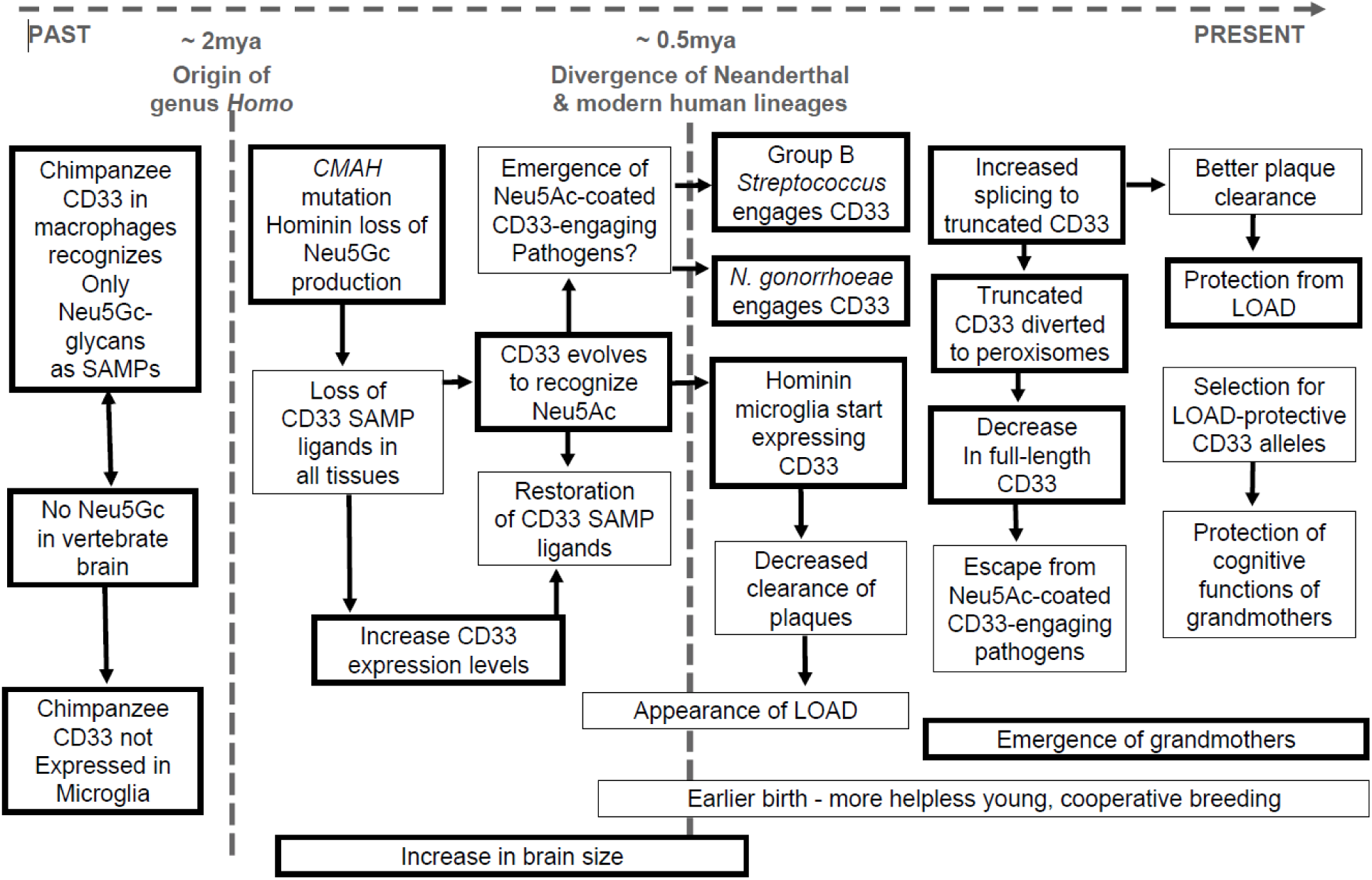
Scenario for evolution of human CD33 in relationship to cell surface sialic acids, infectious disease, brain microglia and cognitive maintenance of grandmothers and other elderly caregivers. This schematic presentation combines the known/likely facts (thick-outlined boxes, including data from this manuscript) as well as suggested possibilities (thin-outlined boxes) into the most likely evolutionary scenario for human-specific evolution of CD33. Starting from the left, the likely chronological order of occurrence is indicated (by arrowheads) with the approximate timeline on the top, along the dotted lines. ‘?’ indicates our reasonable assumption leading to the event. See text for further discussion.

A similar evolutionary scenario appears to underly the case of the human *APOE* gene where variants include both risk alleles (*APOE4)* and protective alleles, (*APOE2*, and *APOE3)* for CVD and LOAD [65]. In this instance, the ancestral *APOE4* allele is associated with increased risks of loss of cognitive functions and the derived alleles may serve to protect the cognition of the elderly caregivers. Interestingly the *APOE4* allele is also correlated with the protection from severe diarrhea in early years of life [66]. Given these examples like APOE and CD33, it remains to be seen how widespread this evolutionary pattern is wherein variants conveying survival advantages in early life coexist with other variants that protect cognition late in life.

## Author Contribution

Experimentation (S.S., N.K., T.C., A.V., A.S., S.D., M.F.), Data analysis (S.S., N.K., T.C., A.V., A.S., S.D., J.M.A., M. F., P. G., A.V.), Critical reagents (H.Y., X.C.), Original draft (S.S., P.G., A.V.), Writing (S.S., N.K., T.C., A.V., A.S., X.C., J.M.A., M.F., P.G., A.V.), Overall supervision (J.M.A., P.G., A.V.), Funding acquisition (A.V.).

## Data Availability

The data for the resampling analysis is available at Code Ocean.

## Declaration of interest

The authors have declared that no conflict of interest exists.

## Methods

### Bacterial culture and cell lines

The bacterial strains used were *Neisseria gonorrhoeae* F62Δ*lgtD* (generous gift from Sanjay Ram, University of Massachusetts Worcester), Group B *Streptococcus* (GBS) strains COH1wt, COH1Δ*neuA*, A909wt and A909Δ*neuA* (generous gifts from Victor Nizet, University of California San Diego). *Neisseria* were grown overnight on chocolate II agar plate and GBS on Todd Hewitt agar plate at 37 ℃ and 5% CO_2_ from the respective frozen glycerol stocks. Prior to the assay, GBS was grown in Todd Hewitt broth at 37 ℃ and 5 % CO_2_ without shaking. The *E. coli* K1 strain was grown in LB. For the CD33 protein purification, HEK293A cells were grown in DMEM media (Invitrogen) containing 10% FCS at 37 ℃ and 5 % CO_2_.

### Sialylation of *Neisseria*

Following overnight growth on chocolate agar plate, the bacteria were grown in GC broth supplemented with IsoVitaleX at 37 ℃, 5% CO_2_ and shaking at 200 rpm in presence or absence of 30 µM CMP-Neu5Ac (Nacalai USA. Inc.) until OD600 equivalent to 0.4– 0.5.

### Bacterial staining

Following appropriate growth, the bacteria were washed with pre-warmed HBSS and stained with 2 µM SYTO13 (Thermo Scientific) for 30 min at 37 ℃ and shaking at 200 rpm in dark. After incubation, the stained bacteria were washed with HBSS and resuspended to a final concentration of OD600 = 1/ml in HBSS for the binding assay.

### Generation of CD33 mutant proteins

A genomic fragment (1228 bp) of human or Chimpanzee CD33(M), including the first 4 exons (2 Ig domains) was fused with pcDNA3.1(-) containing a C-terminal FLAG (EK) sequence followed by a hIgG1-Fc genomic fragment (hinge + 2 Ig-like domains) and described elsewhere [67, 68]. Sixteen mutant variants were made from either construct above using New England Biolabs Q5 site directed mutagenesis Kit according to the manufacturer’s instructions (Supplemental Table 3). Mutagenesis primers listed were designed using NEBaseChanger software.

### Truncated CD33(CD33m) _EK_Fc Construction

U937 cells were cultured in RPMI 1640 supplemented with 10% FCS. Total mRNA was isolated using Qiagens Oligotex Direct mRNA Mini Kit according to the manufacturer’s instructions. CD33m was amplified by PCR using SuperScript III One-Step RT-PCR (Invitrogen) and Gene-specific primers 5’-TTATAT<u>GCTAGC</u>GCCACCATGCCGCTGCTGCTACTGCTGC-3’, NheI site underlined and 5’-GCGCGC<u>GATATC</u>ATGAACCACTCCTGCTCTGGTCTCTTG-3’, EcoRV site underlined. PCR products were run on 2% agarose gel and the 396 bp bands corresponding to CD33(m) were excised and cut with NheI/EcoRV restriction enzymes. Digested bands were sub-cloned into pcDNA3.1(-) containing a C-terminal FLAG (EK) sequence followed by a hIgG1-Fc genomic fragment (hinge + 2 Ig-like domains).

### Purification of CD33 mutants

Transfection supernatants were collected and spun down at 500 g for 5 mins to remove cellular debris. The pH of each supernatant was adjusted to pH 8.0 for optimal binding of protein A-Sepharose beads to hIgG Fc fusion protein. Protein A-Sepharose 4 Fast Flow suspension (GE Healthcare) was washed with Tris-Buffered Saline (TBS) pH 8.0, and a 1:500 ratio of beads:media added to each supernatant. Each tube was subsequently incubated for 24 hrs on a roller in the cold-room. After 24 hours supernatants plus beads were transferred to disposable columns until all liquid has run thru. Beads were washed 3x with TBS pH 8.0 before being eluted directly in 0.3 ml of 1 M Tris-HCl pH 8.0 using 0.1 M Glycine Buffer pH 2.8. Each eluate was put into an Amicon Ultra-15 filter unit with MWCO 30 K for each full length CD33-EK_Fc variant and MWCO 10 K for huCD33m-EK_Fc. Tubes were centrifuged at 4,000 g for 20 mins. Run-through was discarded and the columns washed 3x with TBS pH 8.0. After the last wash, each retentate was recovered from the column and stored at -80°C.

### Binding assay with the bacteria

Bacterial binding with the CD33 proteins were done with the recombinant Fc-chimeric proteins of CD33. Briefly, protein A coated black 96-well plate (Pierce, Thermo Scientific) was washed thrice with TBS containing 0.05% Tween 20 (TBS-T) and coated with 200 ng/well of the respective CD33 protein diluted in 200 mM Tris-HCl pH 8.0, 150 mM NaCl and 1% BSA at 4 ℃ overnight. Following incubation, the coated plate was washed with 200 mM Tris pH 8.0, 150 mM NaCl to eliminate the unbound proteins. Stained bacteria equivalent to OD600 = 0.1 was added to each well of the plate and allowed to interact with the proteins for 30 min at 37 ℃ and 5% CO_2_ without shaking. Following incubation, the plate was washed with TBS-T to eliminate any unbound bacteria and the residual fluorescence was measured upon excitation at 488 nm and emission at 530 nm. The data were analyzed using the excel and Prism software.

### Evolutionary analysis and Detection of positive selection

The protein coding sequences of CD33 were aligned using CLUSTAL W program implemented in MEGA7 and then back translated to obtain a codon alignment. The phylogenetic tree of CD33 protein coding sequences were reconstructed with neighbor-joining method which was implemented in MEGA7 (Figure 1), 1000 bootstrap replicates [69]. The unrooted neighbor joining tree was used for the subsequent analysis.

VCF files were accessed from ftp://ftp.1000genomes.ebi.ac.uk/vol1/ftp/release/20130502/ for 1000 genomes project, http://cdna.eva.mpg.de/neandertal/altai/AltaiNeandertal/VCF/ for Altai Neanderthal and http://cdna.eva.mpg.de/denisova/VCF/hg19_1000g/ for Denisovan. Quality filters were obtained from https://bioinf.eva.mpg.de/altai_minimal_filters/ for Altai and Denisovan. Individuals in 1000 genomes datasets were assigned to populations using http://ftp.1000genomes.ebi.ac.uk/vol1/ftp/technical/working/20130606_sample_info/20130606_sample_info.txt. First, all vcf files were filtered by intersecting with the quality bed files using bedtools intersect (v2.28.0). The filtered vcf files were then combined, per chromosome, to match their position, reference and alternative allele using a custom python script. VCF information was retained along with per-population allele frequencies and archaic genotypes. Next, ancestral alleles were obtained from ensembl (https://rest.ensembl.org/variation/homo_sapiens) by querying each SNP id and appending to the joint vcf entries. The joint vcf files were used as input for further processing in a jupyter notebook to perform resampling analysis. Each SNP of interest is used to select collections of SNPs with matching global allele frequencies, +/- 0.004, or +/- 2 observed haplotypes. A single draw consists of selecting one SNP from each collection to produce a simulated observation and the number of SNPS with derived archaic haplotypes are recorded. After 10,000 such draws, the faction of draws with fewer or equal numbers of derived SNPs is used to produce a p-value estimate. The process is repeated 10,000 times to produce a histogram and provide a confidence estimate on the reported p values (+/- SEM). Methods to replicate the analysis can be found on Code Ocean.

Non-synonymous/ synonymous substitution ratios (ω = dN/dS, or Ka/Ks) have become a useful means for quantifying the impact of natural selection on molecular evolution. In general, the ratio ω = dN/dS is less than one if the gene is undergoing purifying selection, equal to one if the gene is evolving neutrally, and greater than one if positive selection has accelerated the fixation of non-synonymous substitutions that resulted in amino acid changes. The pair-wise computation of Ka/Ks between V-set exon of each species were performed using the program DnaSp v.0 6.0. The initial unrooted tree fed to the program in the format of Newick was: ((Chimpanzee5:0.00000000, Bonobo:0.00000000):0.00222522, Gorilla:0.00979959, Human:0.01351877).

### Molecular Simulation

Starting structures of the V-type domain (residues 18-142) of CD33 were built based on PDB entries 5j0b (A chain) and 6d49 using the graphical interface of YASARA [70]. The two structures differ significantly with respect to the conformation of the C-C’ loop (residues 63-71, compare Supplemental Figure S3). A single mutation (G69R) was introduced into 5j0b to build CD33(human). The initial 3D models of chCD33 were built by swapping residues: N20K, F21I, W22R, A65P, I67V, R69G (in 6d49), L78P, P96L. An N-glycan core (M3) was attached to Asn100. The side chain of His45 was modeled in two conformations (compare Figure 6): ‘down’ (as in PDB entry 6d49) and ‘up’ (as present in PDB entries 5ihb or 5j06 chains A). The systems were solvated in 0.9% NaCl solution (0.15 M) and simulations were performed at 310 K using periodic boundary conditions. The box size was rescaled dynamically to maintain a water density of 0.996 g/ml. Additionally systems were built that contain five molecules of Neu5GcαOMe or Neu5AcαOMe distributed in the simulation box which allowed to simulate binding events. Simulations were performed using YASARA with GPU acceleration [71]. In total 27 MD trajectories were sampled for huCD33 and 20 for chCD33, most of them covering a microsecond timescale (compare Supplemental Figure S4). Conformational Analysis Tools (CAT, http://www.md-simulations.de/CAT/) was used for analysis of trajectory data, general data processing and generation of scientific plots. VMD [72] was used to generate molecular graphics.

### Sialoglycan microarray

The sialoglycan microarray experimental method was adopted from the literature reported earlier [73, 74]. Chemoenzymatically synthesized sialoglycans were quantitated utilizing DMB-HPLC method [75] and 10 mM aqueous stock solutions were prepared. Next, the glycans were diluted to 100 µM in 300 mM Na-phosphate buffer (pH 8.4) and printed in quadruplets on NHS-functionalized glass slides (PolyAn 3D-NHS; catalog# PO-10400401) using an ArrayIt SpotBot® Extreme instrument. The slides were blocked using 0.05M ethanolamine solution in 0.1 M Tris-HCl (pH 9.0), washed with warm Milli-Q water and dried. Printed slides were fitted in a multi-well microarray hybridization cassette (ArrayIt, CA) and rehydrated using 400 µl of ovalbumin (1% w/v, PBS) for one hour in a humid chamber with gentle shaking. The solution was discarded followed by the addition of 400 µl solution of the CD33 protein (30 µg/ml in PBS with 1% w/v ovalbumin) in the individual well. The slides were incubated for 2 h at ambient temperature with gentle shaking followed by washing with PBS-Tween (0.1% v/v) and PBS. The wells were then treated with Cy3-conjugated goat anti-human IgG (1:500 dilution in PBS), incubated for 1h in a dark humid chamber with gentle shaking. After washing and drying, the slides were scanned using a Genepix 4000B scanner (Molecular Devices Corp., Union City, CA) at wavelength 532 nm. Data analysis was performed using the Genepix Pro 7.3 software (Molecular Devices Corp., Union City, CA).

**Supplemental File S1: List of the glycans used for the sialoglycan microarray**. The complete list of the chemoenzymatically synthesized glycans used to determine the binding profile of different CD33 proteins are presented. The binding intensity of the different proteins (indicated on the top of the columns) towards the corresponding glycan are shown in the heatmap (same heatmap as in Figure 3). The red indicates maximum, and blue indicates minimum binding. R = propylamine linker present in the underlying glycan structure. Gal = galactose, GalNAc = *N*- acetylgalactosamine, Glc = glucose, GlcNAc = *N*-acetyl glucosamine, Fuc =L-fucose. The linkage between the monosaccharides is indicated as α- or β- with numbers.

## Acknowledgments

We are very thankful to Dr Sanjay Ram (University of Massachusetts, Worcester) and Dr. Victor Nizet (University of California, San Diego) for the bacterial strains that have greatly benefited the project. This work was supported by NIH grant R01GM32373 and Cure Alzheimer’s Fund grant (to A.V.).

---

**Supplemental Figure S1:**
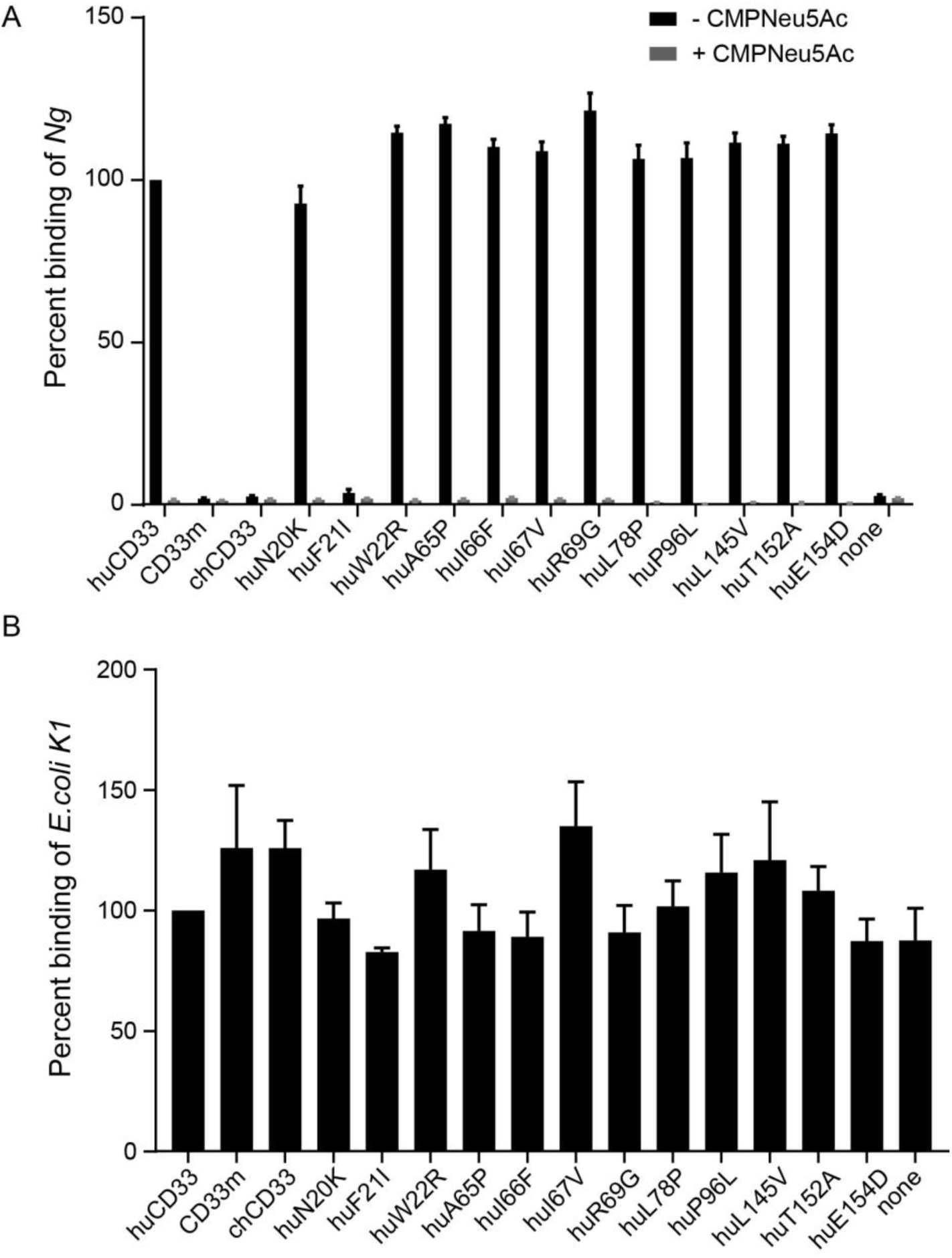
While *E. coli* does not bind CD33, human CD33 binding by *Neisseria* is Sia-dependent. **(A)** Binding of Ng with wildtype or mutant CD33 proteins was determined in the same manner as in Figure 2A. The bacteria for the assay were either grown in presence (+) or absence (-) of exogenous CMP-Neu5Ac as indicated in the legend. All the binding was normalized to wildtype human CD33 binding. Cumulative data from 2 independent experiments, each done in triplet is presented. **(B)** Binding of *E. coli* K1 was determined using the different CD33 proteins. None of the proteins showed any increased binding to the bacteria relative to no protein (control) containing blank well, indicating that there is no binding of the bacteria with the protein.

**Supplemental Figure S2:**
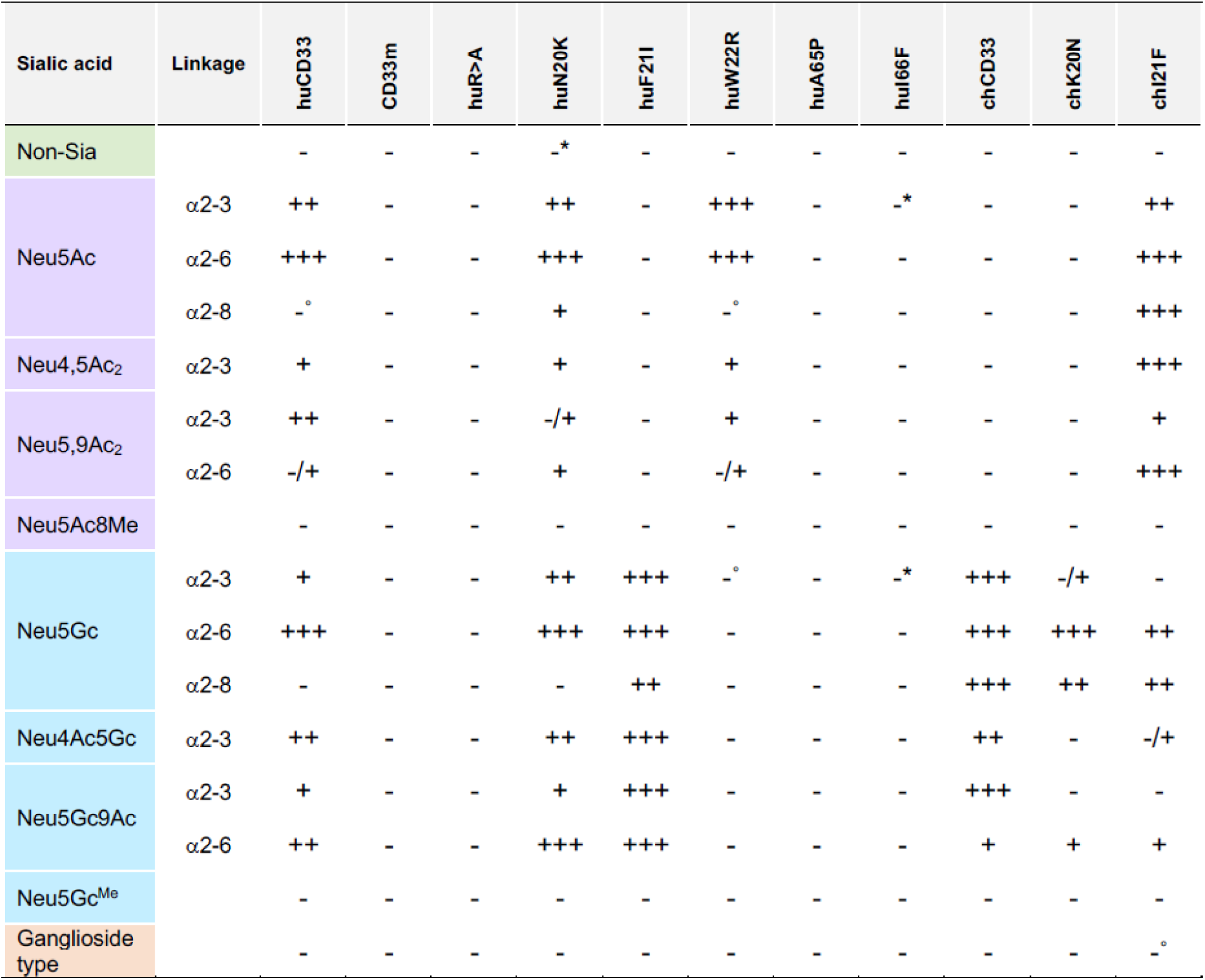
Summarized result of CD33 sialoglycan binding. The results of the sialoglycan microarray binding of different wildtype and mutant CD33 proteins presented in Figure 3 are summarized here. A differential sialoglycan binding preference was observed when wildtype and mutant human/chimpanzee CD33 proteins were tested on the microarray. Binding is annotated with a positive (+) symbol and the strength of the binding is indicated by the number of the symbols. +++ indicates a very strong binding. Negative (-) symbol implies non-binding and -/+ indicates very faint interaction. In some cases, only a few sulfated glycans showed strong binding signal (indicated with asterisk). Degree (°) symbols indicate binding with a very few numbers of glycans only. Linkage indicates the nature of the glycosidic bond of the terminal Sia to the underlying glycan.

**Supplemental Figure S3:**
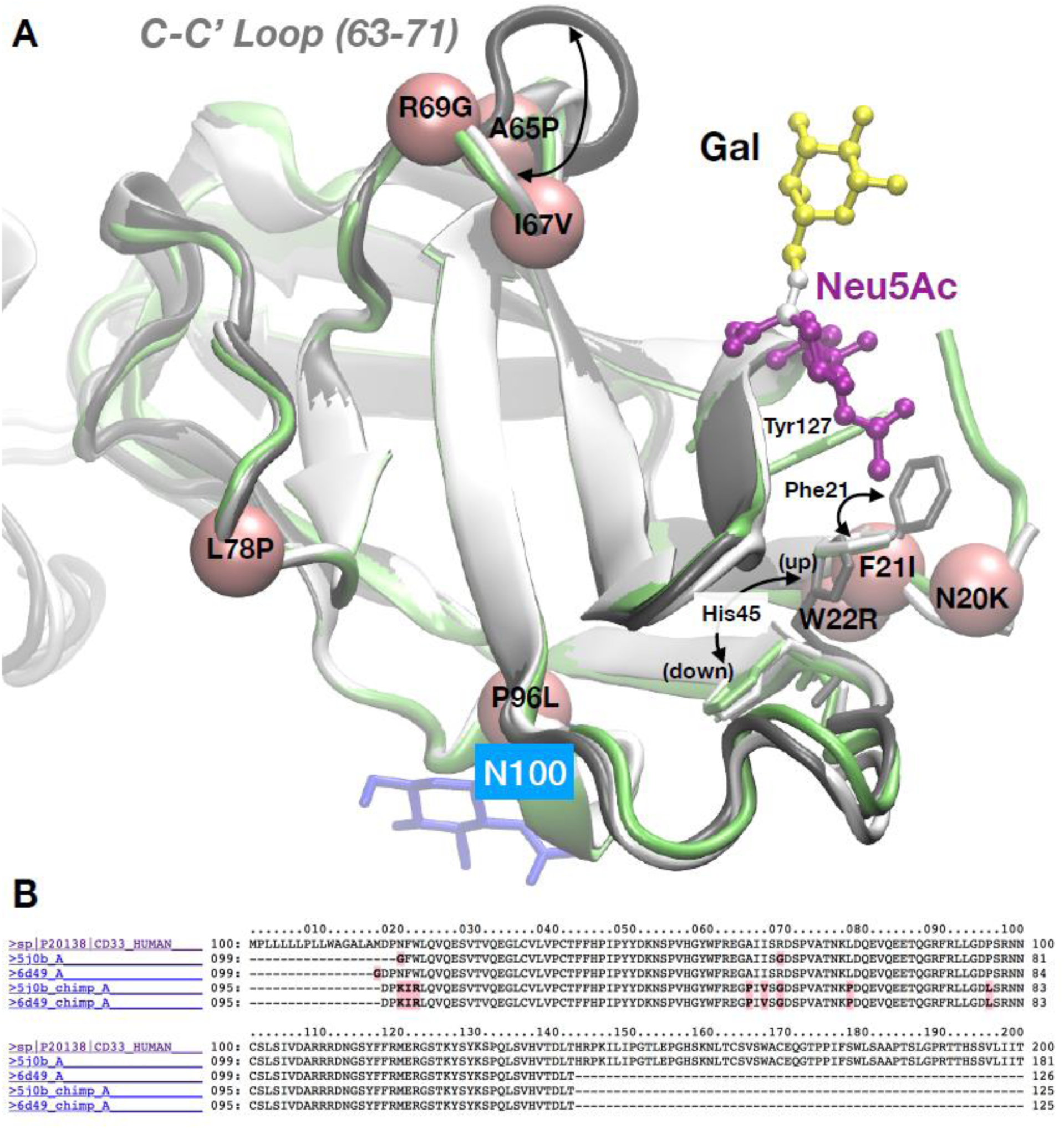
Structure and dynamics of CD33. **(A)** Examples of x-ray structures of huCD33. PDB entries 5ihb (chain A: dark grey, chain B: white), 6d49 (lime). The dynamics of the C-C’ loop and residues Phe21 and His45 are indicated. Positions of mutations present in chimpanzee are labeled on the pink spheres. **(B)** Amino acid sequences. 1: CD33 human (Uniprot), 2-3: PDB entries used for modeling. 3-4: sequences of the chCD33 models.

**Supplemental Figure S4:**
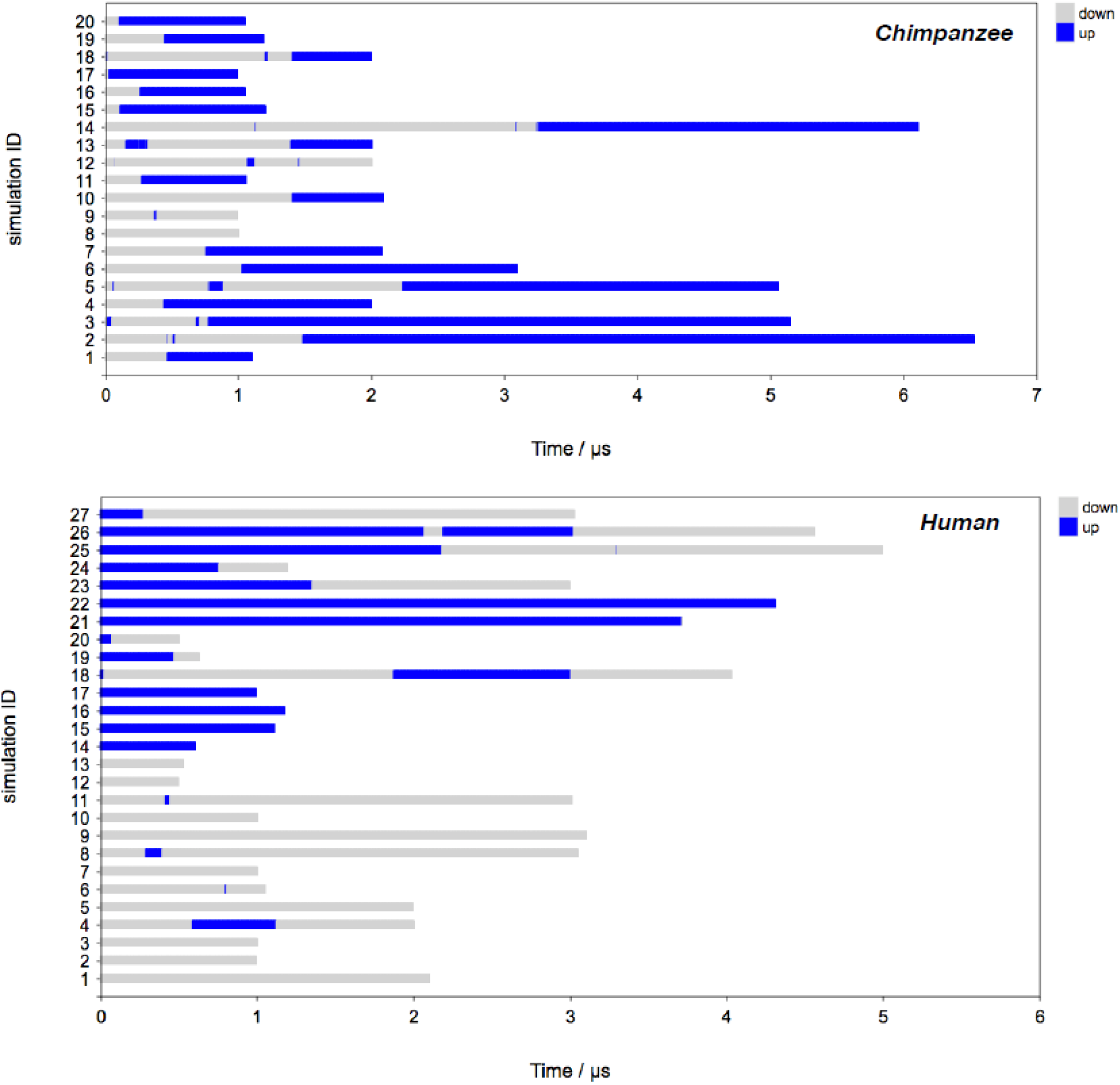
MD trajectories of up/down states of Histidine at position 45 (His 45). Molecular dynamics of His45 side chain orientation. Individual MD trajectories of ‘up’(blue)/‘down’(grey) conformational states are shown. For chCD33, it can be observed reproducibly that simulations started with His45 in a ‘down’ conformational state undergo a transition to the ‘up’ conformational state on the microsecond timescale. Therefore, it may be concluded that chCD33 exists mainly with His45 in an ‘up’ orientation, which would be favorable for binding of Neu5Gc. For huCD33, both conformational states can exist for multiple µs, which explains why huCD33 can bind to Neu5Ac (preferably binds when His45 is ‘down’) and Neu5Gc (preferably binds when His45 is ‘up’).

**Supplemental Figure S5:**
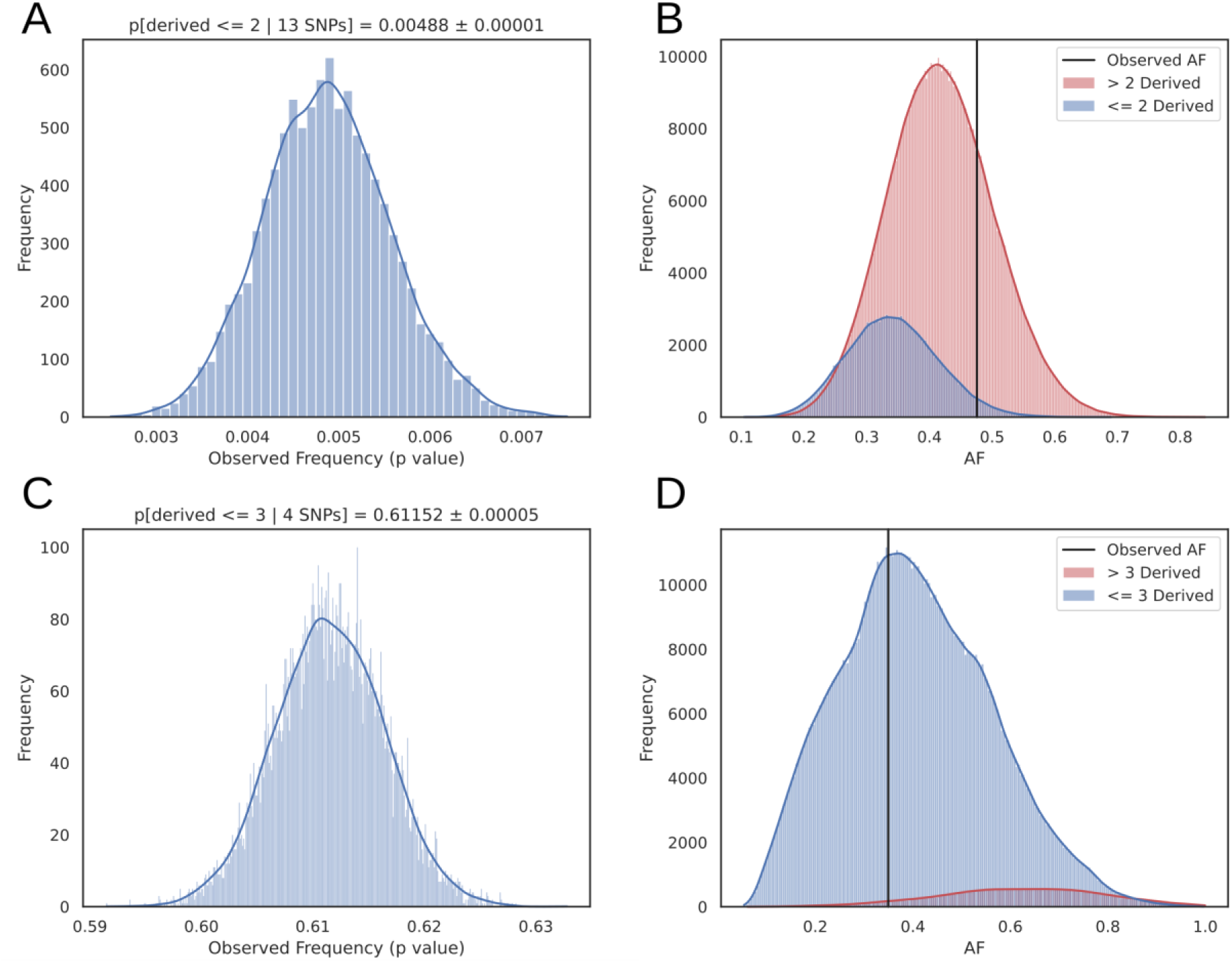
Resampling analysis of the 5.9 million SNPs from 1000 genomes variants. As an alternative to matching AF directly, the set of filtered SNPs were further restricted to those with a derived population frequency greater than 0.05 resulting in a universe of 5.9 million SNPs. We estimated the probability of observing at most two SNPs derived in either the Neanderthal or Denisovan reference genomes and a mean allele frequency as large as the empirical variants of interest (AF = 0.476). By randomly drawing SNPs, we found that the probability of observing 13 SNPs with such as high global allele frequency and lack of derived alleles in archaic genomes to be highly unusual (*p*-value = 0.00487 ± 0.00001) **(A).** The low frequency is driven by two factors, as shown in **(B).** Most of the SNPs sampled have more than two archaic-derived SNPs (red curve). Of those with fewer than two archaic-derived SNPs, the overall allele frequency is typically low compared to the target set. With other Siglec SNPs, resampling captures similar properties **(C** and **D)**, indicating the LOAD protective set does not represent a random sampling from the genome.

**Supplemental Table S1:** Genes affecting cognitive functions in post-reproductive age exhibiting disease-protective alleles uniquely in humans. The corresponding references for each of the genes are mentioned in the table.

| Gene | Associated disease | SNP ID | Description | References |
| --- | --- | --- | --- | --- |
| *CD33* | LOAD | rs12459419,<br>rs3865444 | This study for details | [16, 47] |
| *APOE* | LOAD, CVD | rs7412,<br>rs429358 | Encodes plasma protein APOE, is polymorphic in humans. Three alleles (E2, E3, E4) encode proteins with distinct affinity for lipoprotein particles. The ancestral E4 allele is associated with highest LOAD risk, and increased atherosclerosis and vascular dementia. The derived alleles E2 and E3 seems protective against LOAD, with the lowest risk is in homozygous E2 individuals. | [48, 65, 76] |
| *PICALM* | AD | rs3851179<br>rs10792832 | Encodes phosphatidylinositol-binding clathrin assembly protein (PICALM), considered to be one of numerous reproducible risk genes for LOAD. | [49] |
| *SPON1* | Dementia | rs2618516 | Encodes the developmentally regulated protein F-spondin, reported to be a putative ligand for the amyloid precursor protein (APP). | [50] |
| *TCFLC2* | Diabetes | rs7903146 | Associated with impaired insulin secretion and enhanced hepatic glucose production. | [52] |
| *ARID5B* | AD | rs2588969 | Gene encodes a member of AT-rich interaction domain (ARID) family of DNA binding proteins. The encoded protein forms a histone H3K9Me2 demethylase complex with PHD finger protein 2 and regulates the transcription of target genes involved in adipogenesis and liver development. | [51] |
| *PILRA* | AD | rs1859788 | A cell surface inhibitory receptor that recognizes specific O-glycosylated proteins and expressed on various innate immune cell types including microglia | [53] |
| *CYP3A5* | Salt retention and hypertension | rs776746 | Cytochrome P450 (CYP) genes are abundant in animal, plant, and bacterial genomes and have evolved to metabolize a variety of diverse compounds. | [54] |
| *PPARG* | Diabetes | rs1801282 | A nuclear hormone receptor that regulates adipogenesis | [55] |
| *BIN1* | AD | rs7561528 | Also known as amphiphysin 2, has recently been identified as the most important LOAD risk locus | [20] |
| *SCG2* | Hypertension | rs1017448 | Secretogranin II (SCG2) associates with hypertension | [56] |
| *CAPN10* | Diabetes | rs2975760 | CAPN10 encodes a member of the calpain-like cysteine protease family that regulates blood glucose levels. | [57] |
| *AGT* | Sodium retention | rs699 | Sodium homeostasis links with hypertension | [58] |

LOAD: Late onset Alzheimer's disease; AD: Alzheimer's disease; CVD: Cardiovascular disease

**Supplemental Table S2:** Analysis of Gene variants directly or indirectly affecting cognitive function with their human population frequency. The global frequency of the SNPs identified in Supplemental Table S1 was studied across different populations as indicated in the top of the columns.

| Gene | SNP ID | Allele | Global frequency | African | East Asian | European | South Asian | American |
| --- | --- | --- | --- | --- | --- | --- | --- | --- |
| *CD33* | rs12459419 | C | 0.789 | 0.949 | 0.814 | 0.69 | 0.84 | 0.52 |
|  |  | T | 0.211 | 0.051 | 0.186 | 0.31 | 0.16 | 0.48 |
|  | rs3865444 | C | 0.789 | 0.949 | 0.814 | 0.69 | 0.84 | 0.52 |
|  |  | A | 0.211 | 0.051 | 0.186 | 0.31 | 0.16 | 0.48 |
| *APOE* | rs7412 | C | 0.925 | 0.897 | 0.9 | 0.937 | 0.96 | 0.96 |
|  |  | T | 0.075 | 0.103 | 0.1 | 0.063 | 0.04 | 0.04 |
|  | rs429358 | T | 0.849 | 0.732 | 0.914 | 0.845 | 0.91 | 0.9 |
|  |  | C | 0.151 | 0.268 | 0.086 | 0.155 | 0.09 | 0.1 |
| *PICALM* | rs3851179 | T | 0.351 | 0.105 | 0.407 | 0.371 | 0.39 | 0.39 |
|  |  | C | 0.685 | 0.895 | 0.593 | 0.629 | 0.61 | 0.61 |
|  | rs10792832 | A | 0.313 | 0.094 | 0.409 | 0.372 | 0.4 | 0.39 |
|  |  | G | 0.685 | 0.895 | 0.593 | 0.628 | 0.6 | 0.61 |
| *SPON1* | rs2618516 | T | 0.341 | 0.259 | 0.302 | 0.382 | 0.52 | 0.24 |
|  |  | C | 0.659 | 0.741 | 0.698 | 0.618 | 0.48 | 0.76 |
| *TCFLC2* | rs7903146 | C | 0.772 | 0.74 | 0.977 | 0.683 | 0.7 | 0.77 |
|  |  | T | 0.228 | 0.26 | 0.0023 | 0.317 | 0.3 | 0.23 |
| *ARID5B* | rs2588969 | C | 0.532 | 0.472 | 0.482 | 0.641 | 0.63 | 0.42 |
|  |  | A | 0.468 | 0.528 | 0.518 | 0.359 | 0.37 | 0.58 |
| *PILRA* | rs1859788 | A | 0.341 | 0.102 | 0.612 | 0.321 | 0.29 | 0.5 |
|  |  | G | 0.659 | 0.898 | 0.388 | 0.679 | 0.71 | 0.5 |
| *CYP3A5* | rs776746 | T | 0.379 | 0.82 | 0.287 | 0.05 | 0.33 | 0.2 |
|  |  | C | 0.621 | 0.18 | 0.713 | 0.95 | 0.67 | 0.8 |
| *PPARG* | rs1801282 | C | 0.93 | 0.995 | 0.974 | 0.88 | 0.88 | 0.88 |
|  |  | G | 0.07 | 0.005 | 0.026 | 0.12 | 0.12 | 0.12 |
| *BIN1* | rs7561528 | G | 0.8 | 0.809 | 0.881 | 0.683 | 0.87 | 0.74 |
|  |  | A | 0.2 | 0.191 | 0.119 | 0.317 | 0.13 | 0.26 |
| *SGC2* | rs1017448 | C | 0.879 | 0.635 | 0.963 | 0.979 | 0.97 | 0.95 |
|  |  | T | 0.121 | 0.365 | 0.037 | 0.021 | 0.03 | 0.05 |
| *CAPN10* | rs2975760 | T | 0.882 | 0.971 | 0.907 | 0.841 | 0.79 | 0.87 |
|  |  | C | 0.118 | 0.029 | 0.093 | 0.159 | 0.21 | 0.13 |
| *AGT* | rs699 | A | 0.295 | 0.097 | 0.147 | 0.588 | 0.36 | 0.36 |
|  |  | G | 0.705 | 0.903 | 0.853 | 0.412 | 0.64 | 0.64 |

**Supplemental Table S3:**
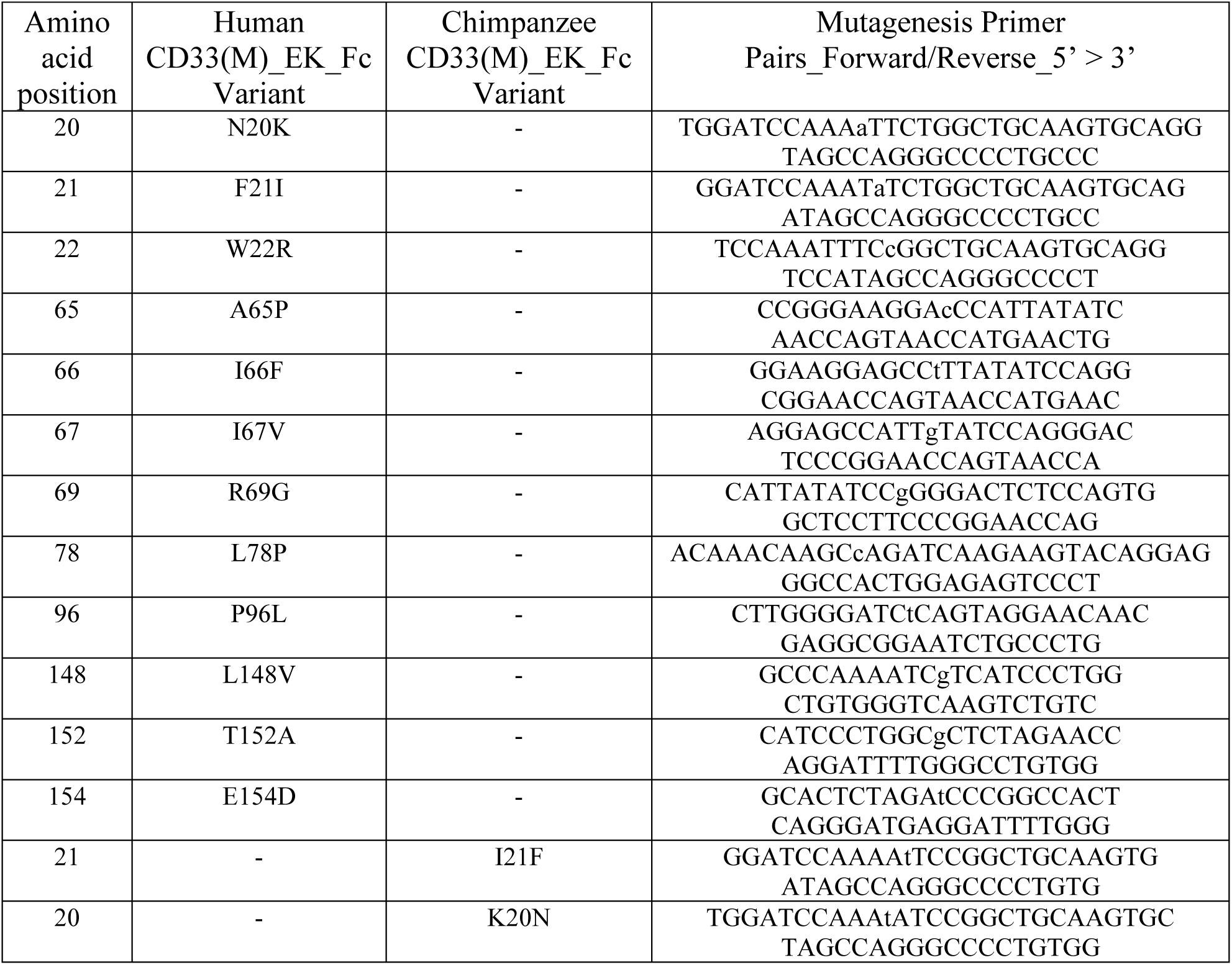
List of the mutagenesis primers used in the study to generate the CD33 mutants. Lowercase letters correspond to base change.

## Notes

### Competing Interest Statement

The authors have declared no competing interest.

